# Glycolysis inhibition functionally reprograms T follicular helper cells and reverses lupus

**DOI:** 10.1101/2024.10.15.618563

**Authors:** Ahmed Elshikha, Yong Ge, Seung Chul Choi, Yuk Pheel Park, Lauren Padilla, Yanan Zhu, William L. Clapp, Eric S. Sobel, Mansour Mohamadzadeh, Laurence Morel

## Abstract

Systemic lupus erythematosus (SLE) is an autoimmune disease in which the production of pathogenic autoantibodies depends on T follicular helper (T_FH_) cells. This study was designed to investigate the mechanisms by which inhibition of glycolysis with 2-deoxy-d-glucose (2DG) reduces the expansion of T_FH_ cells and the associated autoantibody production in lupus-prone mice. Integrated cellular, transcriptomic, epigenetic and metabolic analyses showed that 2DG reversed the enhanced cell expansion and effector functions, as well as mitochondrial and lysosomal defects in lupus T_FH_ cells, which include an increased chaperone-mediated autophagy induced by TLR7 activation. Importantly, adoptive transfer of 2DG-reprogrammed T_FH_ cells protected lupus-prone mice from disease progression. Orthologs of genes responsive to 2DG in murine lupus T_FH_ cells were overexpressed in the T_FH_ cells of SLE patients, suggesting a therapeutic potential of targeting glycolysis to eliminate aberrant T_FH_ cells and curb the production of autoantibodies inducing tissue damage.

## Introduction

High-affinity class-switched autoantibodies, the main pathogenic effectors in SLE, are crucially regulated by T follicular helper (T_FH_) cells, regardless of whether they differentiate through the germinal center (GC) or the extrafollicular route^1^. The frequency of T_FH_ cells correlates with disease activity in SLE patients^2,3^, and pharmacological inhibition of IL-21, a cytokine produced at high levels by T_FH_ cells, ameliorated disease in lupus-prone mice^4^. T_FH_ cells from the triple congenic (TC) B6.*Sle1.Sle2.Sle3* lupus mice present activated transcriptomic effector programs, including increased T cell receptor (TCR) signaling as well as cytokine and co-receptor expression, as compared to T_FH_ cells from B6 congenic healthy controls^5^. In addition, naive TC CD4^+^ T cells (T_N_) are poised to differentiate into T_FH_ cells. These results indicated that lupus genetic susceptibility skews CD4^+^ T cells toward differentiation into highly active T_FH_ cells.

T cell metabolism is altered in SLE patients and mice, and thus, some of these alterations may point to novel therapeutic targets^6^. We have shown that the combined inhibition of glycolysis and oxidative phosphorylation reverse disease in multiple models of SLE in correlation with decreased CD4^+^ T cells activation^7^. T_FH_ cells are sustained by both glycolysis and oxidative phosphorylation (OXPHOS)^8,9^. Inhibiting glycolysis through HK2 with 2DG^10^ or PKM2 with TEPP-46^11^, impaired T_FH_ cell polarization in a cell-intrinsic manner. The inhibition of glycolysis *in vivo* with 2DG or TEPP-46 showed that the expansion of T_FH_ cells and the resulting production of autoantibodies is highly dependent on glucose in several strains of lupus-prone mice^11,12^. In contrast, T_FH_ cell expansion induced by immunization with a foreign antigen or influenza infection was not affected by 2DG^12^. Thus, these results suggest that autoreactive lupus T_FH_ cells may depend more on glycolysis than T_FH_ cells induced by foreign antigens.

In the present study, we elucidated mechanisms by which the inhibition of glycolysis reduces the frequency and reprograms the functions of T_FH_ cells in lupus. We used the (NZW x BXSB.Yaa)F1 mouse model of the disease, herein referred to as W.Yaa, in which the Yaa duplication of the *Tlr7* gene largely controls pathogenesis through TLR7/type I interferon (IFN) signaling^13^, a pathway critically in SLE^14^. 2DG treatment completely reversed the autoimmune manifestations and renal pathology in W.Yaa mice^15^. The W.Yaa model is thus highly relevant to elucidate the mechanisms leading to the rewiring of cellular glycolysis to potentially control autoreactive T_FH_ cells and thus mitigate disease progression. Here, we demonstrate that 2DG reprograms T_FH_ cell activation and metabolism and normalizes the corresponding gene expression and DNA methylation. Particularly, the inhibition of glycolysis restored mitochondrial homeostasis as well as autophagolysosomal flux, which was critically impaired by type I IFN-enhanced chaperone-mediated autophagy (CMA). The functional reprogramming of lupus T_FH_ cells by 2DG was demonstrated by their protective effect upon adoptive transfers. Finally, the transcript signatures affected by 2DG in T_FH_ cells from W.Yaa mice overlap with the signatures of T_FH_ cells from SLE patients. Thus, these results clearly underline that uncontrolled glycolysis drives dysfunctional T_FH_ cells and its inhibition reprograms the mitochondrial and autophagolysosomal machinery in these cells to attenuate disease progression.

## Results

### Glycolysis controls T_FH_ cell expansion in W.Yaa mice

We confirmed here (Figure S1A - C) that a 2DG treatment initiated in anti-dsDNA IgG positive W.Yaa mice prevented the development of fatal renal pathology and reversed autoantibody production^15^. 2DG reduced the frequency of W.Yaa T_FH_ cells as well as the frequency of follicular regulatory (T_FR_) cells to levels similar to B6 healthy controls. The effect was stronger on T_FH_ cells, thereby reducing the high T_FH_ to T_FR_ ratio that has been associated with SLE^16^ (Figure 1A - D). Extrafollicular helper T (T_EXFH_) cells provide critical help to lupus extrafollicular B cells to produce autoantibodies^17^. W.Yaa mice develop a robust T_EXFH_ cell expansion, which was also reduced by 2DG (Figure 1D). Consistent with a reduced frequency, 2DG decreased the proliferation of T_FH_, T_FR_ and T_EXFH_ cells (Figure S1D). 2DG also reduced ICOS expression on T_FH_ and T_EXFH_ cells, as well as in T_N_ cells from W.Yaa mice and T_FH_ cells from B6 mice (Figure 1E). Since ICOS induces *Bcl6* expression at the early stage of T_FH_ cell differentiation and maintains their phenotype^18^, a higher ICOS expression contributes to T_FH_ cell expansion. TCR activation triggers *Icos* transcription^19^. Consistent with a stronger TCR signaling in lupus T_FH_ cells, *Icos* message expression was higher in W.Yaa than in B6 T_FH_ cells, and it was decreased by 2DG (Figure 1F). On the other hand, ICOS protein level is negatively regulated through ubiquitination by CBL/CBL-b, which are expressed at lower levels in the CD4^+^ T cells from SLE patients^20^. W.Yaa T_FH_ cells expressed lower levels of *Cbl* message than W.Yaa T_N_ and B6 T_FH_ cells, but it was not changed by 2DG (Figure S1E). At the protein level, CBL and CBL-b were expressed at lower levels in CD44^+^CD4^+^ T cells (surrogates for T_FH_ cells whose numbers were highly reduced by 2DG) than T_N_ cells, but at a similar level between B6 and W.Yaa cells. However, 2DG increased CBL and CBL-b expression in W.Yaa CD44^+^CD4^+^ T cells (Figure 1 G, H). These results suggest that 2DG reduced ICOS expression both at the transcriptional level downstream and at the protein level through CBL/CBL-b.

**Figure 1.**
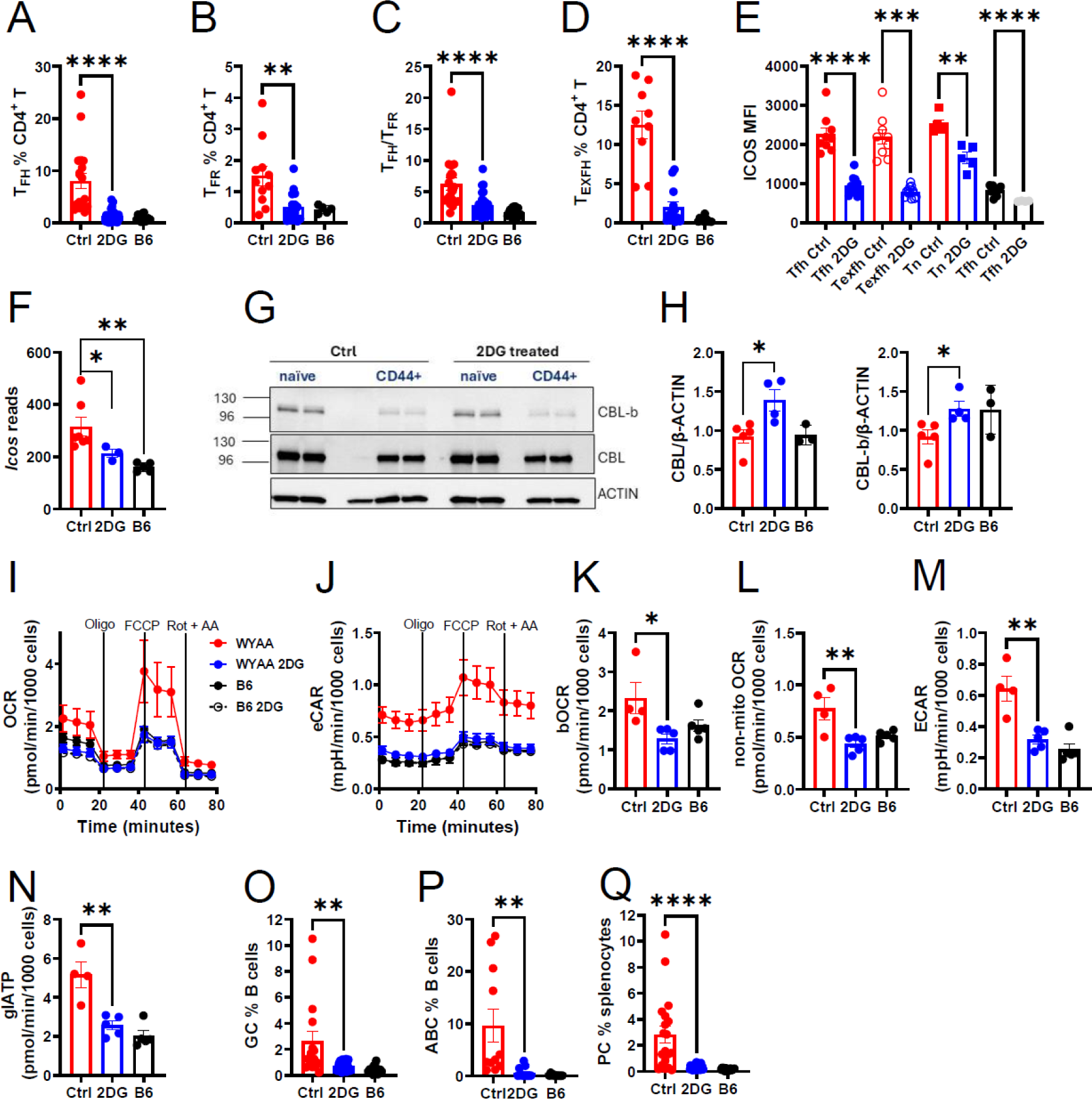
2DG treatment reduced the expansion of T cells and B cells involved in autoantibody production in W.Yaa mice. Frequency of T_FH_ (**A**) and T_FR_ (**B**) cells in total CD4^+^ T cells. (**C**) T_FH_/T_RF_ ratio. (**D**) Frequency of T_EXFH_ cells in total CD4^+^ T cells. (**E**) ICOS MFI in T_FH_, T_FR_ and T_EXFH_ cells. (**F**) *Icos* expression in T_FH_ cells (RNAseq results, N = 3 – 7). **(G** – **H)** Western blot analysis of CBL and CBL-b expression with representative image of T_N_ and CD44^+^CD4^+^ T cells from W.Yaa mice treated with 2DG or control (**G**) and quantitation in CD44^+^CD4^+^ T cells from B6 and W.Yaa mice (**H**) (N = 3 – 5). Oxygen consumption rate (OCR) (**I**) and extra-cellular acidification rate (ECAR) (**J**) in a mitochondrial stress assay in CD44^+^ CD4^+^ T cells. Vertical lines indicate the injection of oligomycin (oligo), FCCP and rotenone + antimycin A (rot +ant). Basal OCR (**K**), non-mitochondrial OCR (**L**), basal ECAR (**M**) and glycolytic ATP production (**N**). Frequency of GC B (**O**) and ABC (**P**) cells in total B cells, and frequency of plasma cells (PC) in splenocytes (**Q**). 1-way ANOVA with multiple comparisons tests in which the difference between Ctrl and B6 was highly significant (in A – E and L – Q, not shown). Mean + SEM, *: P < 0.05; **: P < 0.01; ***: P < 0.001; ****: P < 0.0001. W.Yaa mice treated with 2DG and controls, and age-matched B6 mice (A – E and O – Q: N = 10 – 21; I – M: N = 4 - 5).

W.Yaa CD44^+^CD4^+^ T cells showed enhanced respiration, glycolysis, non-mitochondrial respiration, as well as ATP generated from glycolysis, all of which were reduced by 2DG to B6 levels (Figure 1I - N). Notably, 2DG treatment had no effect on these parameters in B6 CD44^+^CD4^+^ T cells. 2DG drastically reduced the frequency of GC B cells, age-related Tbet^+^ B cells (ABCs), and plasma cells (Figure 1O - Q). However, respiration and glycolysis were not affected by 2DG in W.Yaa B cells (Figure S1H), suggesting that the effect of 2DG on B cell differentiation may be secondary to its effect on CD4^+^ T cells. Overall, these results demonstrate that the inhibition of glycolysis curtailed the expansion of T_FH_ cells as well as related T_FR_ and T_EXFH_ cells in lupus-prone W.Yaa mice, in association with a reduced ICOS expression and a normalization of glycolysis and respiration.

### Glycolysis alters TCR signaling and mitochondria homeostasis in lupus T_FH_ cells

Next, we focused on the effect of 2DG on activation and metabolic markers in W.Yaa T_FH_ cells. We evaluated strain and cell-specific differences in T_FH_ cells relative to T_N_ cells in both W.Yaa and B6 mice. We then compared these markers between T_FH_ cells from W.Yaa mice treated with 2DG and untreated controls, with B6 T_FH_ cells as a reference. First, we examined TLR7 expression, which corresponds to *Trl7* gene duplication in the W.Yaa model. TLR7 activation as well as type 1 IFN, its downstream target, inhibit T_FH_ cell differentiation^21,22^. However, the T_FH_ cell population expands despite high type I IFN activity in SLE patients and in lupus models with the Yaa-TLR7 duplication^23^ (and this report). As expected, TLR7 expression was higher in W.Yaa than B6 T cells, but only W.Yaa T_FH_ cells expressed higher levels of TLR7 than T_N_ cells, which were reduced by 2DG (Figure 2A). This suggests that increased TLR7 signaling contributes to W.Yaa T_FH_ cell expansion in a glucose-dependent manner. TCR-dependent activation of RelA/p65 increases cMyc expession, which triggers glycolysis^24^. TCRβ expression was lower on T_FH_ than on T_N_ cells in both strains, which may correspond to the exhausted/anergic phenotype of this T cell subset^25^, and it was not altered by 2DG (Figure 2B). Consistent with a lower TCRβ expression, B6 T_FH_ cells downregulated RelA and cMyc compared to T_N_ cells (Figure 2C - D). However, W.Yaa T_N_ cells expressed higher levels of RelA and cMyc, which were further increased in W.Yaa T_FH_ cells, and reduced by 2DG. These results, thus, suggest a stronger TCR signaling in W.Yaa T_FH_ cells, potentially supporting glycolysis, as well as *Icos* expression.

**Figure 2.**
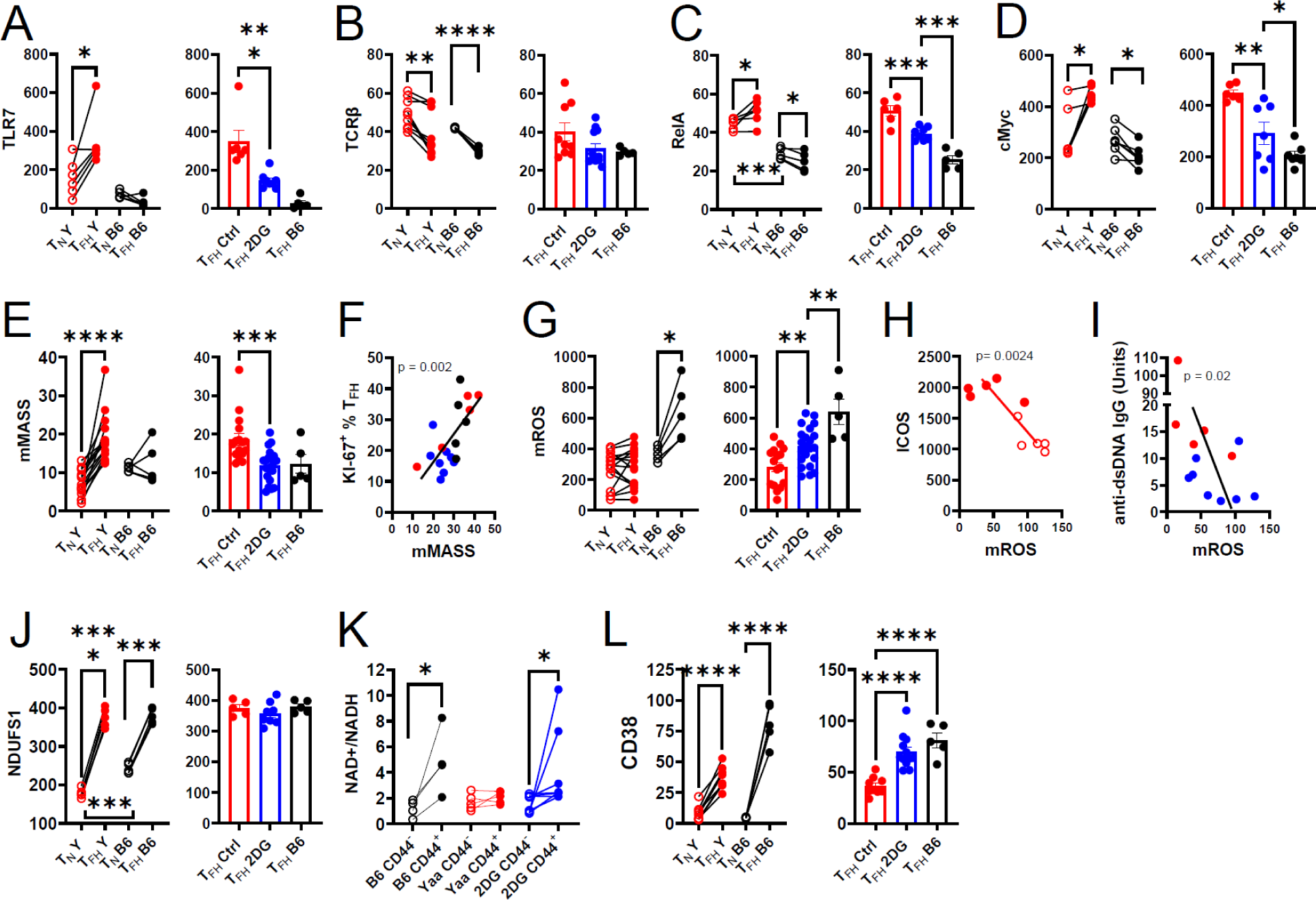
Glycolysis alters TCR signaling and mitochondria homeostasis in lupus T_FH_ cells. (**A** – **E, G, J** and **L**) Graphs on the left show T_N_ and T_FH_ cells from untreated W.Yaa (Y) and B6 mice compared with paired *t* or Wilcoxon matched-pair signed rank tests between T_N_ and T_FH_ cells in the same mice, and by *t* tests comparing T_N_ between strains. Graphs on the right compare the same parameters in T_FH_ cells between untreated controls (Ctrl) and 2DG-treated (N= 6 – 19) W.Yaa mice, and age-matched untreated B6 (N = 5). Comparisons were made with Dunnett’s T3 multiple comparisons tests. (**A**) TLR7; (**B**) TCRβ; (**C**) RelA p65; (**D**) cMyc; (**E**) Mitochondrial mass; (**G**) Mitochondrial ROS. Correlations between mMASS and the frequency of KI-67^+^ in mice in T_FH_ cells from the 3 groups (**F**), mROS and ICOS on untreated W.Yaa T_N_ and T_FH_ cells (**H**) or serum anti-dsDNA IgG in treated and untreated W.Yaa mice (**I**). (**J**) NDUFS1 expression. (**K**) NAD^+^/NADH ratio in treated and untreated W.Yaa and B6 CD44^+^CD4^+^ and CD44^-^CD4^+^ T cells from the same mice. (**L**) CD38 expression. Correlation statistics were obtained with Pearson or Spearman tests, depending on the normality of the data distribution. All values, except the NAD^+^/NADH ratio, are shown as mean fluorescence intensity (MFI) measured by flow cytometry. Mean + SEM, *: P < 0.05; **: P < 0.01; ***: P < 0.001; ****: P < 0.0001.

In contrast to B6, W.Yaa T_FH_ cells increased their mitochondrial mass (mMASS) compared to T_N_ cells (Figure 2E), with a strong correlation between mMASS and T_FH_ cell proliferation (Figure 2F). B6 T_FH_ cells increased mROS production relative to their T_N_ cells (Figure 2G), which is consistent with the high oxidative state of non-autoimmune T_FH_ cells^26^. This was not the case for W.Yaa T_FH_ cells despite their increased mMASS. 2DG normalized mMASS and partially restored mROS levels in W.Yaa T_FH_ cells (Figure 2E and G). Reduced mROS correlated strongly with ICOS expression in CD4^+^ T cells from untreated W.Yaa mice as well as with serum anti-dsDNA IgG in the entire W.Yaa cohort (Figure 2H, I). We next investigated electron transport chain complex I, a major source of mROS^27^. The expression of NDUFS1, a key complex I subunit, strongly increased in T_FH_ compared to T_N_ cells in both strains, but started from a lower level in W.Yaa T_N_ cells (Figure 2J). 2DG had no effect on NDUFS1 expression. As expected from increased complex I levels in T_FH_ cells, the NAD^+^/NADH ratio increased in B6 CD44^+^CD4^+^ T cells, a surrogate for T_FH_ cells to meet cell number requirements for the assay, compared to T_N_ cells. However, this was not the case for W.Yaa CD44^+^CD4^+^ T cells unless they were from 2DG-treated mice (Figure 2K). Relatively reduced NAD^+^ level might be due to increased expression of the NAD^+^ degrading enzyme CD38, which is overexpressed in CD4^+^ T cells from SLE patients^28^. CD38 expression increased in T_FH_ compared to T_N_ cells in both strains, consistent with T cell activation upregulating CD38 (Figure 2L). However, CD38 expression was lower on W.Yaa T_FH_ than on B6 T_FH_ cells, suggesting that their lower NAD^+^ level is unlikely due to increased processing by CD38. Interestingly, 2DG treatment restored CD38 expression on W.Yaa T_FH_ cells to B6 levels (Figure 2L). A reduced synthesis could also decrease NAD^+^ level in W.Yaa T_FH_ cells. NAMPT, the rate-limiting enzyme in the NAD^+^ salvage pathway, is overexpressed in CD4^+^ T cells from SLE patients and lupus-prone MRL/*lpr* mice^29^. *Nampt* expression was also higher in W.Yaa T_FH_ cells than in B6 (FDR = 0.0015), but it was not altered by 2DG, suggesting a glycolysis-independent increased NAD^+^ biosynthesis in W.Yaa T_FH_ cells. Overall, these results suggest that defective mitochondrial functions in W.Yaa T_FH_ cells decrease the NAD^+^/NADH ratio and reduce mROS production, which correlates with ICOS expression and autoantibody production. Furthermore, these changes were mitigated by inhibiting glycolysis with 2DG.

T_FH_ cell differentiation requires both mTORC1 and mTORC2 activation, at least in part through glycolysis^8^. Accordingly, both W.Yaa and B6 T_FH_ cells showed an increased expression of pEBP1 and pS6 (mTORC1 targets) and pAKT Ser473 (mTORC2 target) (Figure S1I - K). However, mTORC1 activation was similar in W.Yaa and B6 T_FH_ cells and was minimally affected by 2DG. pAKT Ser473 level was lower in W.Yaa than in B6 T_FH_ cells and unaffected by 2DG. These results, thus, suggest that the glucose-dependent expansion of T_FH_ cells in W.Yaa mice may not depend on mTOR activation.

### Inhibiting glycolysis normalized the transcriptome of W.Yaa T_FH_ cells

To elucidate the mechanisms potentially involved in the glycolysis-dependent expansion of T_FH_ cells in W.Yaa mice, we first performed bulk RNA-seq on FACS-sorted T_N_ and T_FH_ cells from W.Yaa mice compared to T_N_ and T_FH_ cells from B6 controls, respectively. For both transcriptomic and metabolomic analyses, T_FH_ cells were sorted as CD4^+^CD44^+^PD-1^+^PSGL1^lo^ cells, a validated gating strategy that maximizes cell viability by not including intracellular stains for CXCR5 and FOXP3^30^, which also includes T_FR_ and T_EXFH_ cells. We have shown that gene expression overlaps extensively between CD4^+^CD44^+^PD-1^+^PSGL1^lo^ and CD4^+^PD1^+^CXCR5^+^ cells^5^. Moreover, consistent with the effect of 2DG on individual T_FH_, T_FR_ and T_EXFH_ subsets (Figure 1A - D), 2DG decreased the frequency of CD4^+^CD44^+^PD-1^+^PSGL1^lo^ cells and increased the frequency of T_N_ cells in W.Yaa mice, while it had no effect on the frequency of these subsets in B6 mice (Figure S1L, M). W.Yaa T_FH_ cells showed a distinct gene expression signature compared to B6 T_FH_ cells, with a large number of differentially expressed genes (DEGs) in both directions (Figure S2A, B). Top upregulated genes in W.Yaa T_FH_ cells included *Fcnb*, which encodes for Ficolin-1, a disease biomarker in SLE patients^31^ as well as *Eno1b*, a retrotransposon-encoded homolog of the gene encoding for *enolase*, a key glycolytic enzyme^32^. The expression of the glucose and lactate transporters, *Glut1* and *Slc16a3*, as well as most glycolytic enzymes, was higher in W.Yaa T_FH_ cells (Figure S3), supporting the observed higher glycolytic activity. Cell cycle and cytokine-cytokine receptor interaction were the two main signaling pathways enriched in W.Yaa T_FH_ cells, consistent with high proliferation, differentiation and effector functions (Figure S2C, D). The Lysosome pathway was also overexpressed in W.Yaa T_FH_ cells, including *Lamp2* as well as cathepsin genes such as *Ctsd*, encoding for cathepsin D, the major lysosomal aspartic protease, and *Ctse*, encoding for cathepsin E, which is expressed at high level in pathogenic CD4^+^ T cells in lupus-prone MRL/*lpr* mice^33^. As reported in TC mice^5^, the majority of DEGs between W.Yaa and B6 T_FH_ cells were shared with their respective T_N_ cells (Figure S2E). Indeed, gene expression in T_N_ cells was different between the two strains, including the lysosome and several metabolic pathways (Figure S4). T_FH_-specific DEGs were dominated by the cytokine-cytokine receptor interaction pathway (Figure S2F). These results combined with those obtained in the TC lupus-prone model^5^ suggest that a lupus-associated genetic program promoting differentiation and proliferation T_FH_ cells initiates in T_N_ cells, with additional T_FH_-specific programs enhancing their effector functions.

Next, we compared the transcriptomic profiles of T_FH_ cells between W.Yaa mice treated with 2DG and untreated controls. The inhibition of glycolysis markedly altered gene expression in W.Yaa T_FH_ cells (Figure 3A), including the reduced *Il21* level (Figure 3B) that is consistent with a reduced T_FH_ function. At the pathway level, 2DG reduced the lysosome, cytokine-cytokine receptor interaction and cell cycle pathways (Figure 3C, D), which are pathways differentially expressed between W.Yaa and B6 T_FH_ cells (Figure S2C, D). The cytokine pathway responded to 2DG by the inhibition of T_FH_ and T_H1_-related genes (e.g., *Il21, Il10, Il12r1, Il12rb2*, *Ifng*, *Ifngr)* combined with increased expression of T_H2_-related cytokines and cytokine receptors (e.g., *Il4, Il13ra1, Il9r*). In addition, 2DG reversed the overexpression of glycolytic genes (Figure S3) combined with an upregulation of the OXPHOS pathway, with an enrichment of mitochondrial (e.g., *mt-Nd2, mt-Co2*, *mt-Cytb*) and chromosomally encoded respiratory genes (e.g., *Nadufa5, Atp5e*, and *Atp5k*). The extensive DEG overlap between W.Yaa and B6 T_FH_ cells on one hand and between 2DG-treated vs. control W.Yaa T_FH_ cells on the other hand suggests that W.Yaa T_FH_ cells were transcriptionally reprogrammed by 2DG.

**Figure 3.**
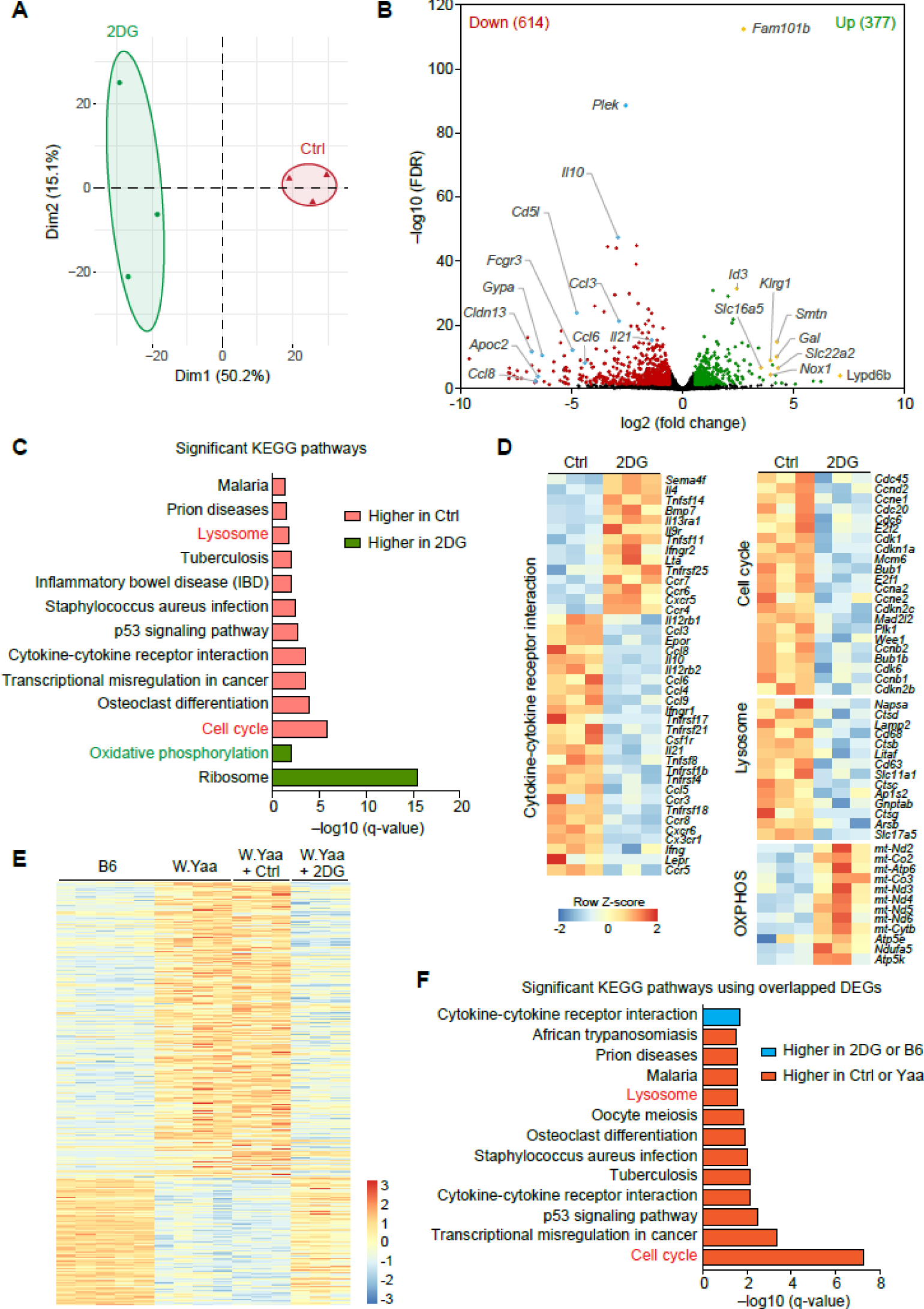
The inhibition of glycolysis normalized the W.Yaa T_FH_ transcriptome. RNA-seq was performed on T_FH_ cells from 2DG-treated or untreated controls (Ctrl) W.Yaa mice (N = 3 per group). (**A**) Principal component analysis. (**B**) Volcano plot of DEGs up-regulated (green) or downregulated (red) in 2DG-treated W.Yaa T_FH_ cells. (**C**) KEGG pathways differentially enriched (green: up, red: down) in 2DG-treated W.Yaa T_FH_ cells. (**D**) Heat maps of DEGs in the Lysosome, Cytokine-cytokine receptor interaction, Cell cycle and Oxidative phosphorylation pathways. (**E**) Combined heatmap of T_FH_ DEGs between B6 vs. W.Yaa (N = 4 per group) and between W.Yaa control vs. 2DG-treated (N = 3) W.Yaa mice, showing that 2DG restored a B6-like signature in W.Yaa T_FH_ cells with significantly overlapping KEGG pathways (**F**).

### Inhibiting glycolysis promotes DNA methylation in W.Yaa T_FH_ cells

DNA hypomethylation in immune-related genes has been widely reported in SLE patients and correlates with disease activity^34^. A genome-wide analysis showed that CpG methylation was higher in T_FH_ cells from 2DG-treated W.Yaa mice as compared to untreated controls. Further analysis defined 357 hyper- and 75 hypo-differentially methylated regions (DMRs) in 2DG-treated T_FH_ cells (Figure 4A), 14.6% of which were in promoters and enhancers (Figure 4B). A correlation analysis identified 16 DEGs with altered CpG methylation within their promoters, enhancers or introns, 13 of which had both a reduced expression and increased DNA methylation in T_FH_ cells from the 2DG-treated group (Figure 4C). These included *Ifng* and *Il10,* both of which are involved in lupus pathogenesis^35,36^, as well as *Lag3,* whose expression in T_FH_ cells has been associated with IL-10 or IL-21 production in pathogenic settings^37,38^. T_FH_ signature genes *Irf4*, *Ikzf3* and *Blimp1* were also downregulated and hypermethylated. Relevant to the observed mitochondrial alterations in W.Yaa T_FH_ cells that were responsive to 2DG, *Irf4* regulates mitochondrial homeostasis in plasma cells^39^ and IKAROS (encoded by *Ikzf3*) has been linked to mitochondrial metabolism and glycolysis in macrophages^40^. Other genes with hypomethylated DMRs include *Mid1,* which links mitochondrial respiration to calcineurin during autophagy^41^, and *Vmp1*, which is critically implicated in autophagosome formation^42^, both parts of the lysosome pathway. *Ubash3a* modulates TCR signaling^43^, and *Kcnk5* has been linked to calcineurin signaling, and it regulates the phosphorylation of numerous mitochondrial proteins^44^. Examination of specific DMRs in 4 of these genes confirmed the increased methylation levels in T_FH_ cells from 2DG-treated mice (Figure 4D). Overall, these results suggest that the inhibition of glycolysis reverts DNA hypomethylation that may control the expression of genes contributing to the W.Yaa T_FH_ cell pathogenic signature.

**Figure 4.**
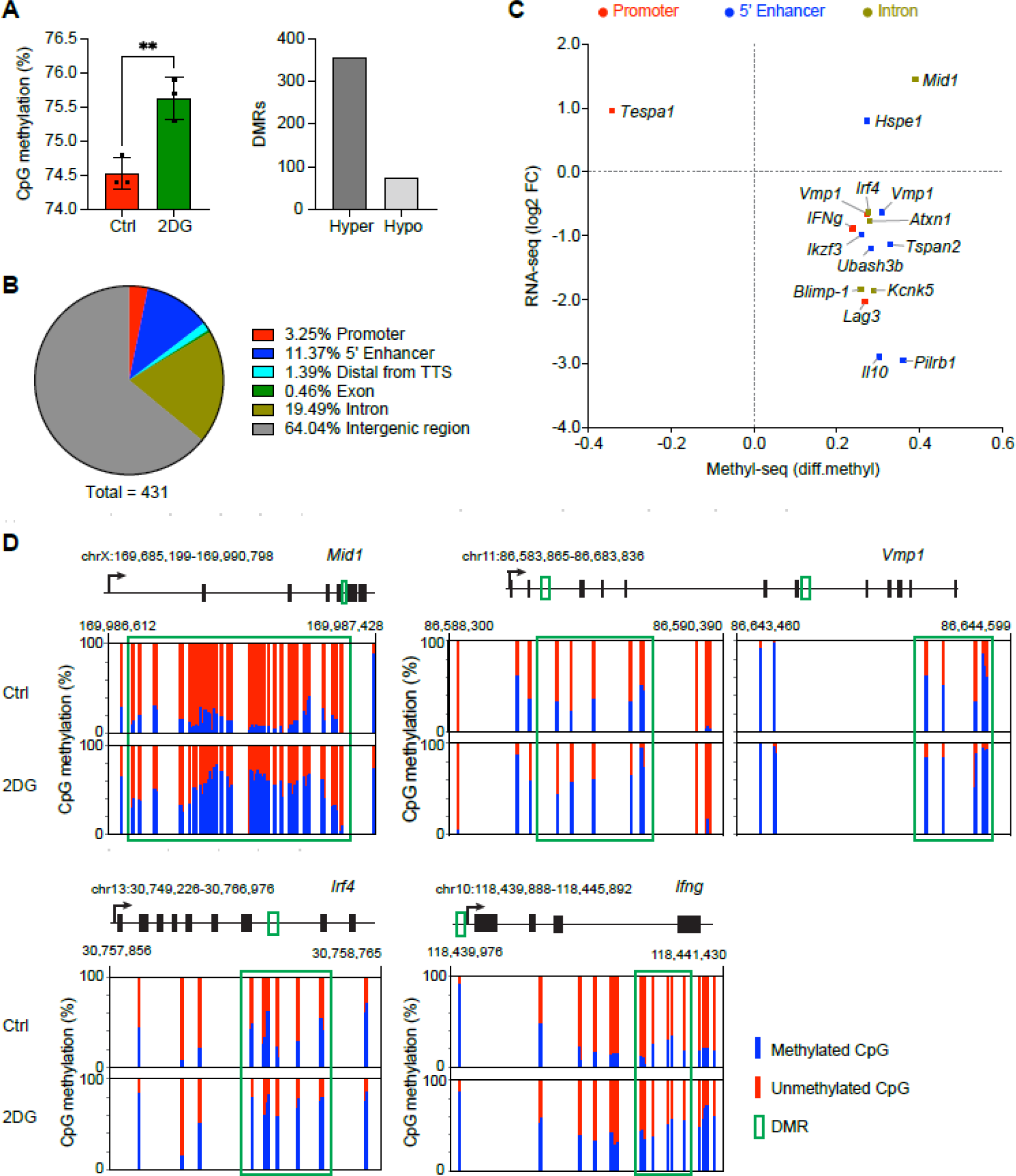
The inhibition of glycolysis increased DNA methylation in W.Yaa T_FH_ cells. (**A**) Whole genome CpG methylation ratios in T_FH_ cells from W.Yaa mice treated with 2DG or controls (N = 3 each), with distribution of hyper-or hypomethylated DMRs in the 2DG-treated samples shown on the right. Mean + SEM compared with a *t* test. **: P < 0.01. (**B**) Distribution of DMRs relative to protein-coding genes. (**C**) DEGs (Y axis) with differential CpG methylation (X axis) in their promoter (red), enhancer (blue), or intron (green), showing 13/16 them with combined increased methylation and decreased expression. (**D**) DMRs in 4 genes, *Mid1*, *Vmp1*, *Irf4* and *Ifng*, shown in panel C, comparing the distribution of methylated and unmethylated CpGs between T_FH_ cells from 2DG-treated and control W.Yaa mice.

### Inhibiting glycolysis normalized metabolites in W.Yaa T_FH_ cells

Untargeted metabolomic analysis of T_FH_ cells isolated from W.Yaa and B6 mice demonstrated altered sialic acid metabolism, pentose phosphate pathway (PPP), as well as ascorbate and aldarate metabolism (Figure S5A), a key carbohydrate metabolic pathway that protects cells from oxidative damage. An increased ribulose and a decreased gluconate levels indicate an activated oxidative PPP flux relative to the gluconate shunt, as observed in the immune cells of SLE patients^45^. The PPP supports nucleotide synthesis, here shown as guanine and cytosine, in agreement with enhanced the cell cycle pathway found in the transcriptome of W.Yaa T_FH_ cells (Figure 3C, D). 2-oxoadipate and 5-acetamidopentanoate belong to tryptophan metabolism, which is altered in SLE and has been associated with T cell activation in lupus-prone mice^46,47^. In addition, anti-inflammatory and antioxidant glycolate and N-acetylneuraminate^48,49^ were reduced in W.Yaa T_FH_ cells. 2DG altered most of these pathways, and the individual metabolites showed similar abundances in T_FH_ cells from B6 and 2DG-treated W.Yaa mice compared to W.Yaa controls (Figure S5B, C). Mitochondrial metabolites (e.g., carnitine, succinate) were enriched by 2DG in T_FH_ cells (Figure S5B), consistent with 2DG increasing the OXPHOS transcriptomic signature (Figure 3C, D) and improving mitochondrial functions (Figure 2).

### Inhibiting glycolysis normalized autophagolysosomal defects in W. Yaa T_FH_ cells

Conflicting lysosome and autophagy defects have been reported in different immune cell types in SLE patients and murine models of the disease^50^. Thus we investigated the lysosomal gene signature expressed by W.Yaa T_FH_ cells and normalized by 2DG. The size of the lysosome as well as its activity, measured with Lysotracker-Red and DQ-Red BSA quenching, respectively, increased in T_FH_ cells compared to T_N_ cells in both mouse strains, but to a greater extent in W.Yaa T_FH_ cells, and both were reduced by 2DG (Figure 5A, B). However, the expression of TFEB, the transcription factor responsible for lysosome biogenesis, was similar between W.Yaa and B6 T_FH_ cells (Figure S6A). The autophagy vacuole content was larger in both T_N_ and T_FH_ cells from W.Yaa mice as compared to B6 mice, in which it moderately increased in T_FH_ relative to T_N_ cells (Figure 5C). Moreover, the 2DG treatment reduced the autophagic vacuoles from W.Yaa T_FH_ cells to B6 levels. The size of the autophagic vacuole content as well as the amount of LC3, an autophagosome marker, increased with age and thus with the development of autoimmunity in W.Yaa T_FH_ cells (Figure S6B). Consistent with an expanded lysosome, the expression of lysosomal marker LAMP1 was higher in T_FH_ cells than in T_N_ cells. However, LAMP1 expression was lower in W.Yaa than in B6 T_FH_ cells, and it was partially rescued by 2DG (Figure 5D). LAMP2, a mediator of autophagolysosomal fusion, also increased in T_FH_ cells compared to T_N_ cells but, contrary to LAMP1, it was reduced by 2DG (Figure 5E). The resulting skewed LAMP2/LAMP1 ratio in W.Yaa T_FH_ cells compared to B6 T_FH_ cells was restored by 2DG (Figure 5F). Accordingly, W.Yaa CD4^+^CD44^+^ T cells showed large LAMP1^+^ areas with unfused LC3 autophagosomes that were normalized in T cells from 2DG-treated mice, which displayed colocalized and reduced LAMP1 and LC3 staining (Figure 5G, S6C). These results suggest that the inhibition of glycolysis rescued an impaired autophagolysosome clearance in W.Yaa T_FH_ cells.

**Figure 5.**
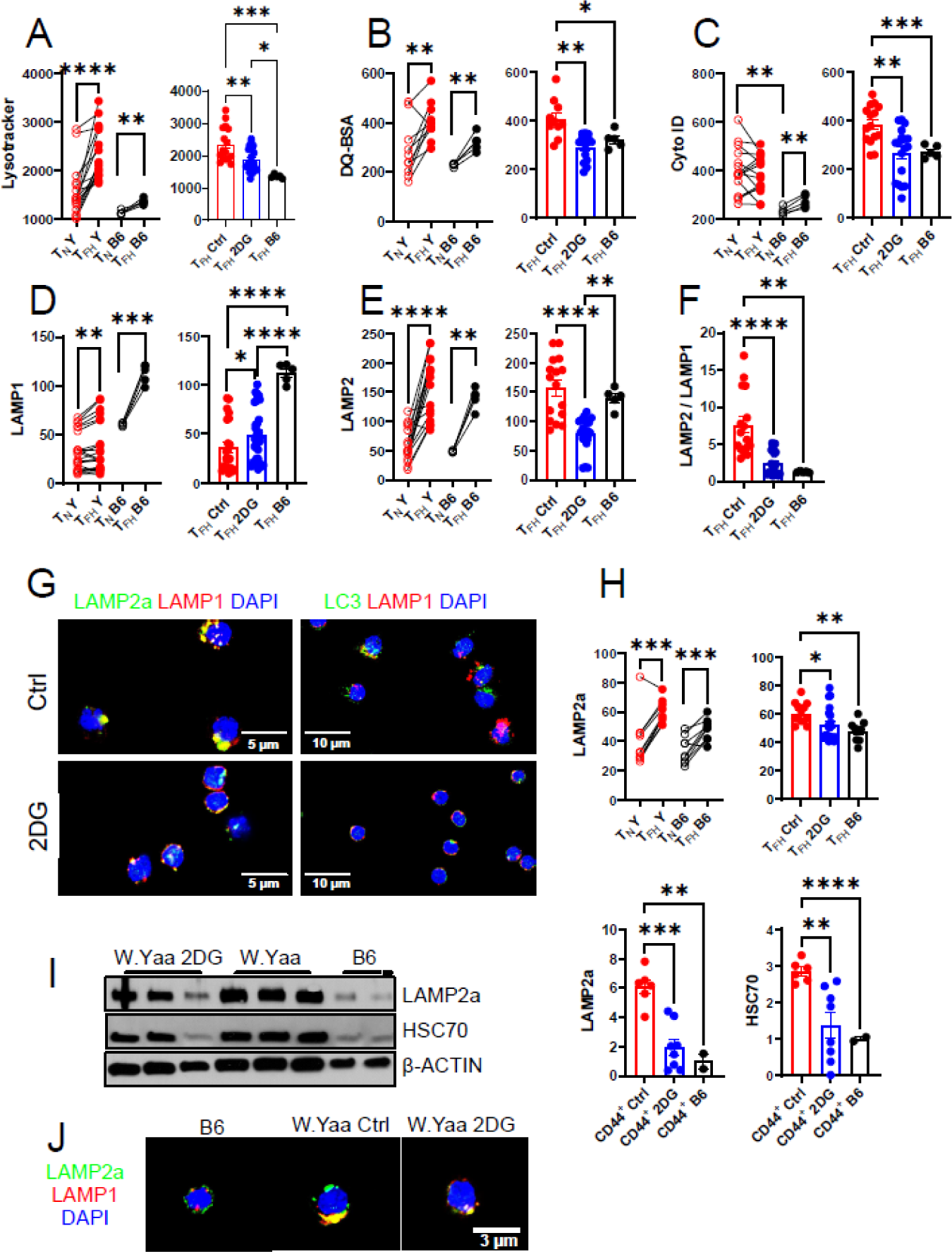
The inhibition of glycolysis reduced chaperone-mediated autophagy in W.Yaa T_FH_ cells. (**A** – **F**). Graphs on the left show T_N_ and T_FH_ cells from untreated W.Yaa (Y) and B6 mice with paired *t* or Wilcoxon matched-pair signed rank tests comparing T_N_ and T_FH_ cells in the same mice, and by *t* tests comparing T_N_ between strains. Graphs on the right compare the same parameters in T_FH_ cells between untreated control (Ctrl) and 2DG-treated (N= 14 - 34) W.Yaa mice, and age-matched untreated B6 (N = 5). Comparisons were made with Dunnett’s T3 multiple comparisons tests. (**A**) Lysotracker; (**B**) DQ-BSA; (**C**) Cyto-ID; (**D**) LAMP1; (**E**) LAMP2; **F**. LAMP2 / LAMP1 ratio. **G**. Representative confocal images of CD4^+^CD44^+^ T cells from 2DG-treated and control W.Yaa mice stained with LAMP2a, LAMP1 and DAPI (left) and with LC3, LAMP1 and DAPI (right). (**H**) LAMP2a as in A - E, shown as MFI measured by flow cytometry. **I**. Representative Western blot analysis of CD4^+^CD44^+^ T cells from 2DG-treated, control W.Yaa and B6 mice probed for LAMP2a and HSC70 with quantification relative to β-ACTIN shown on the right. Dunnett’s T3 multiple comparisons tests. (**J**) CD4^+^CD44^+^ T cells from B6, W.Yaa control and 2DG-treated stained with LAMP2a, LAMP1 and DAPI at a higher magnification. Mean + SEM, *: P < 0.05; **: P < 0.01; ***: P < 0.001; ****: P < 0.0001.

Although 2DG decreased LC3 levels in W.Yaa T_FH_ cells, it had no effect on the classical markers of macroautophagy (Figure S6D, E). Moreover, activation-induced macroautophagy was not altered by 2DG in isolated CD4^+^ T cells (Figure S6F). We investigated whether the high LAMP2 expression in W.Yaa T_FH_ cells might correspond to LAMP2a, a splice isoform that mediates CMA. LAMP2a expression increased in T_FH_ cells compared to T_N_ cells in both mouse strains, but it was higher in W.Yaa than in B6 T_FH_ cells and reduced by 2DG (Figure 5H, I). The expression of CMA chaperone HSC70 in T_FH_ cells mirrored that of LAMP2a (Figure 5I). Accordingly, CD4^+^CD44^+^ and CD4^+^ W.Yaa T cells accumulated LAMP2A^+^ LAMP1^+^ lysosome structures, which were reduced by 2DG (Figure 5G, J and S6C).

We then investigated whether TLR7 signaling could be responsible for the induction of glycolysis-dependent CMA in W.Yaa T_FH_ cells. The 2DG treatment reduced the expression of type I IFN-stimulated genes (ISG) in W.Yaa CD4^+^ T cells (Figure S7A), suggesting a reciprocal relationship between TLR7-induced type I IFN and glycolysis. Next, we validated the results obtained in W.Yaa T_FH_ cells using the related B6.*Sle1.*Yaa strain, in which disease development is slower^51^. B6.*Sle1*.Yaa mice showed a higher frequency of T_FH_ cells than B6.*Sle1* mice, and both strains showed a higher frequency of T_FH_ cells than B6 mice (Figure S7B). LAMP2a expression increased in B6.*Sle1.Yaa* T_FH_ cells compared to B6 and B6.*Sle1* cells, but not in their T_N_ cells (Figure S7C, D). To directly test that TLR7 activation induced LAMP2a expression in T_FH_ cells, we treated B6, B6.*Sle1* and TC mice with the TLR7/8 agonist R848 (Figure S7E), which increased their T_FH_ cell frequency (Figure S7F, H). Interestingly, both T_N_ and T_FH_ cells showed a higher expression of LAMP2a in R848-treated mice than in the corresponding untreated control mice (Figure S7G, I). These results suggest that TLR7 signaling and most likely its downstream product type I IFN increased glycolysis-dependent CMA in T_FH_ cells.

HK2, one of the three rate-limiting enzymes of glycolysis and the target of 2DG, is selectively degraded by macroautophagy^52^. Thus, defective autophagic flux is thought to increase HK2 levels and potentiate glycolysis. Accordingly, HK2 expression was increased in W.Yaa CD4^+^ T cells in parallel with LC3 expression, and reduced by 2DG (Figure S7). In the absence of CBL and CBL-b, the resulting high level of ICOS signaling prevented the CMA-mediated degradation of BCL-6 expression through the PI3K/AKT pathway^20^. Here, we showed that BCL-6 expression was similar between W.Yaa and B6 T_FH_ cells, and it was not affected by 2DG. There was no correlation between ICOS and BCL-6 expression in these T_FH_ cells (Figure S1F, G). In addition, pATK Ser473 level was lower in W.Yaa than in B6 T_FH_ cells and unaffected by 2DG (Figure S1K). Finally, CMA was increased in in W.Yaa relative to B6 T_FH_ cells and reduced by 2DG. Therefore, there is no evidence in this spontaneous model of SLE that high ICOS expression prevents BCL-6 degradation, or that the reduction of T_FH_ cell expansion by 2DG is mediated through this pathway.

Overall, our results suggest that W.Yaa T_FH_ cells demonstrate an altered autophagolysosomal flux associated with increased CMA induced by TLR7 activation. Inhibition of glycolysis by 2DG restored the macroautophagic flux and reduced CMA in T_FH_ cells while reducing their frequency, suggesting that TLR7-induced T_FH_ cell expansion and defective autophagolysosomal machinery may be associated with uncontrolled glycolysis in these cells.

### 2DG-reprogramed T_FH_ cells protected mice against lupus nephritis

To test whether 2DG functionally reprogrammed T_FH_ cells and could mitigate disease progression in lupus-prone mice, we performed adoptive transfers of T_FH_ cells sorted from 2DG-treated or control W.Yaa mice into pre-autoimmune mice with the same background (Figure 6A). Additionally, age-matched W.Yaa control mice received PBS. Transfers of W.Yaa 2DG-T_FH_ cells reduced the relative amount of anti-dsDNA IgG (Figure 6B) and ameliorated renal pathology as evidenced by lower T cell and macrophage infiltrates, smaller glomeruli, and complement C3 deposits (Figure 6C - E). Furthermore, there was a trend toward lower GN scores and elimination of tubulointerstitial lesions (Figure 6F - H), which are associated with poor SLE outcomes^53^. We validated these results in the TC lupus model. Transfers of TC 2DG-T_FH_ cells reduced splenocyte numbers and anti-dsDNA IgG (Figure 6I, J), and alleviated renal pathology with smaller glomeruli, fewer C3 deposits, and reduced T cell and macrophage infiltration (Figure 6K - M). Combined results from the two mouse strains showed a marked reduction in serum anti-dsDNA IgG, lower frequencies of splenic T_FH_ and GC T cells, and reduced mTORC1 activation in CD4^+^ T cells (Figure 6N - Q). Since 2DG reduced the T_FR_ cell frequency (Figure 1B) co-transferred with T_FH_ cells, it is unlikely that T_FR_ cells were responsible for the improved outcomes of 2DG-T_FH_ recipients. Overall, these results suggest that inhibition of glycolysis by 2DG functionally rewires T_FH_ cells toward an immunomodulatory phenotype, resulting in ameliorating lupus-like pathology in the hosts.

**Figure 6.**
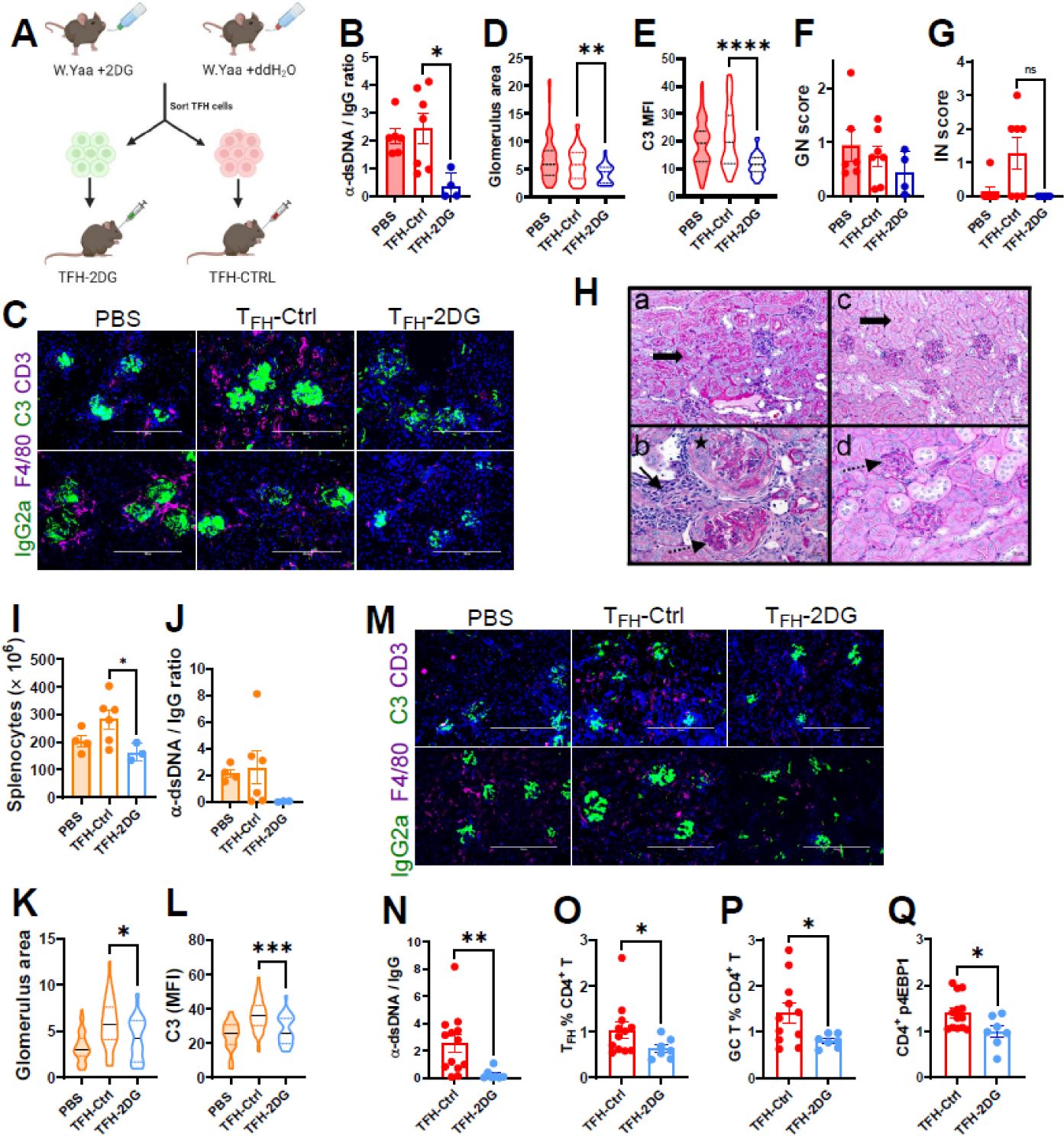
T_FH_ cells from 2DG-treated mice protected from the development of lupus-nephritis. (**A**) T_FH_ cells from 2DG-treated or control lupus-prone mice were transferred into young mice from the same strain and compared to age-matched mice that received PBS. (**B – H**). 6 weeks old W.Yaa mice were evaluated 8 weeks after T_FH_ cell transfer (N = 6 PBS, 7 Ctrl, and 4 2DG from 3 cohorts). (**B**) Anti-dsDNA IgG / IgG ratio. (**C**) Representative kidney sections stained with antibodies against C3 and CD3 (top) and IgG2a and F4/80 (bottom). Quantitation of the glomerulus surface area (**D**) and C3 deposit (**E**) (PBS: 60, Ctrl: 83 and 2DG: 34 glomeruli). Glomerulonephritis (**F**) and interstitial (**G**) scores evaluated on PAS-stained kidney sections with representative shown images in (**H**): **a** and **b**: Ctrl, **c** and **d**: 2DG (a and c: 200x; b and d: 400x). In **a**, arrow points to prominent proximal tubular droplets, and shows normal proximal tubules in **c**. In **b**, the star shows a fibrocellular crescent, the plain arrow an immune infiltrate, and the dashed arrowpoints to glomerular endocapillary hypercellularity and mesangial expansion. In **d**, the dashed arrow points to a mild mesangial expansion with normal glomerular cellularity. (**I** – **M**) 8 weeks old TC mice were evaluated 5 months after T_FH_ cell transfer (N = 4 PBS, 6 Ctrl and 3 2DG). (**I**) Splenocytes numbers. (**J**) Anti-dsDNA IgG / IgG ratio. Quantitation of the glomerulus surface area (**K**) and C3 deposits (**L**) evaluated by immunofluorescence, with representative kidney sections shown in (**M**). (**N** – **Q**) Combined results between the two strains. (**N**) Anti-dsDNA IgG / IgG ratio. Frequency of splenic T_FH_ (**O**) and GC T (**P**) cells. (**Q**) Normalized p4EBP-1 expression in CD4^+^ T cells. **B**, **D** – **G** and **I** – **L**: Dunnett’s T3 multiple comparisons tests. **N** – **Q**: *t* tests. **D** – **E**, **K** – **L**: violin plots with medians and quartiles. All other graphs: Mean + SEM. *: P < 0.05; **: P < 0.01; ***: P < 0.001; ****: P < 0.0001.

### T_FH_ cells from SLE patients and W.Yaa mice share a transcriptome partially controlled by glycolysis

As previously reported^2,3^, the frequency of peripheral T_FH_ cells was higher in SLE patients than in healthy controls (HCs); however, there was no difference in T_PH_ cell frequency (Figure S9A, B). Bulk RNA sequencing showed different transcriptomic profiles between T_FH_ cells from SLE patients and HCs (Figure 7A), resulting in 504 up- and 151 down-regulated genes (Figure 7B). Lysosomal-associated membrane protein (*LAMP3*) was one of the most upregulated genes. Importantly, overexpression of *LAMP3* in the salivary glands of patients with Sjögren’s syndrome was associated with lysosomal dysfunction and impaired autophagy^54^. Other overexpressed genes in SLE T_FH_ cells were *MAFB*, encoding for transcription factor MAF-BZIP, which is part of the T_FH_ cell lineage program^55^, as well as *ALOX5*, which is strongly activated by mitochondrial dysfunction and lipid peroxidation^56^. Importantly, intracellular iron accumulation, which leads to lipid peroxidation, drives the expansion of lupus T_FH_ cells^57,58^. Finally, anti-apoptotic *BCL2* was overexpressed in SLE T_FH_ cells, which we have found to be also overexpressed in murine lupus T_FH_ cells^5^, suggesting that apoptosis resistance might sustain the expansion of SLE T_FH_ cells. Gene set enrichment analysis showed that SLE T_FH_ cells were dominated by type I IFN signaling (Figure 7C). Furthermore, SLE-T_FH_ cells showed enriched “positive regulation of B cell proliferation”, suggesting that they may possess an enhanced effector function. Interestingly, the mitochondrial electron transport pathway was also enhanced in SLE T_FH_ cells (Figure 7C, D), which might be a compensatory expression due to defective mitochondrial metabolism. This is supported by a strong TCA cycle signature in the metabolites of T_FH_ cells from SLE patients, which contained reduced levels of key TCA intermediates, including succinate, isocitrate and fumarate (Figure S9D, E).

**Figure 7.**
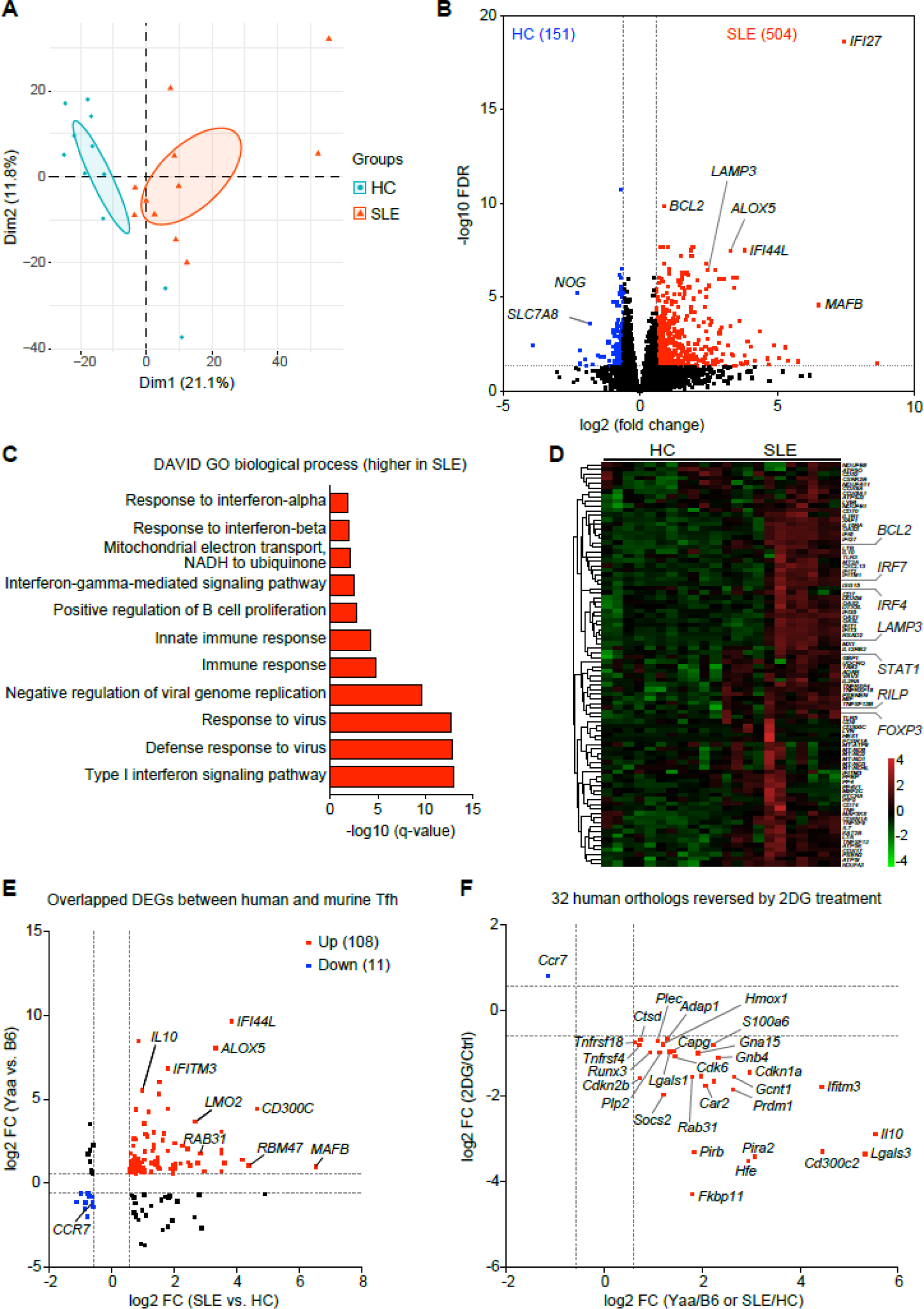
T_FH_ cells from SLE patients shared a 2DG-sensitive gene signature with the W.Yaa T_FH_ cells. (**A**) PCA analysis of the transcriptome of T_FH_ cells isolated from SLE patients and healthy controls (HCs). (**B**) Volcano plot of DEGs up-regulated in SLE patients (red) or HC (blue) T_FH_ cells. (**C** - **D**) Significant pathways enriched in SLE compared to HC T_FH_ cells with corresponding heatmap. (**E**) Overlap of human and murine ortholog DEG between SLE vs. HC (X axis) and W.Yaa vs. B6 (Y axis) T_FH_ cells. (**F**) Overlap of human and murine ortholog DEGs in T_FH_ cells from both human and mouse lupus vs. controls (X-axis) and from 2DG-treated vs. control W.Yaa mice (Y axis).

Finally, we identified 108 up- and 11 down-regulated murine orthologs shared between the SLE patient / HC T_FH_ DEGs and the W.Yaa / B6 DEGs T_FH_ DEGs (Figure S2), which included *ALOX5, MAFB* and *IL10* (Figure 7E). ToF increase clinical relevance, we then correlated the 119 overlapping DEGs with the W.Yaa T_FH_ genes that responded to 2DG (Figure 3). Obtained results suggest that 32 human orthologs overexpressed in the T_FH_ cells of SLE patients can be reversed by inhibiting glycolysis (Figure 7F). These include genes involved in the cell cycle (*CDK6, CDKN1A, CDKN2B*), B cell signaling (*LILRB4A*, *PIRB*, *PIRA2*), cytokine signaling (*Il10, SOCS2*), iron hemostasis (*HMOX1, HFE*) and galectins (*LGALS1, LGALS3*), which regulate cell activation and inflammation, highlighting the role that glycolysis may play in expanding T_FH_ cells in SLE patients.

## Discussion

SLE develops from a combination of genetic susceptibility and environmental risk factors through a complex pathogenesis with many components that remain unclear. There is consensus, however, that the loss of immune tolerance leading to the production of autoantibodies against nucleic acid/protein complexes is at the root of the disease. In this process, the expansion of autoreactive B cells that differentiate into autoantibody-secreting cells is tightly controlled by T_FH_ cells^59^. Accordingly, expansion of T_FH_ cells has been consistently observed in patients with SLE and in lupus-prone mice^1,60,61^, although the underlying mechanisms are largely unknown. We have reported that spontaneous T_FH_ cells from several lupus-prone mouse models are uniquely sensitive to the inhibition of glycolysis, suggesting that targeting glucose metabolism could selectively reduce autoantibody production^12^. Here we investigated the mechanisms by which 2DG restrains the expansion of lupus T_FH_ cells in W.Yaa mice, a model of TLR7-driven lupus, in which 2DG completely reversed clinical disease^15^. 2DG is a glucose analog that competitively blocks glucose phosphorylation by HK2 at the first step of glycolysis. Treating W.Yaa mice with 2DG reduced T_FH_ cell frequency to B6 control levels. Treatment with a drug that prevents the translocation of PKM2, another glycolytic enzyme, to the nucleus where it induces the expression of glycolytic genes, also reduced the T_FH_ cell frequency in another model of lupus^11^, confirming that inhibiting glycolysis targets lupus T_FH_ cells. Reduction of T_FH_ cell frequency by 2DG was associated with a reduced frequency of both GC B cells and ABCs, as well as the autoantibodies they produce, delaying clinical disease^12^ (and this study). However, 2DG normalized the elevated glycolysis of W.Yaa T_FH_ cells, but not B cells, suggesting that T_FH_ cells were the direct targets of the treatment. We thus focused on T_FH_ cells from 2DG-treated W.Yaa mice compared to heathy controls. However, we cannot rule out that the inhibition of glycolysis also impacted W.Yaa B cells, as we have previously shown in the TC lupus model^62^.

We showed here that, compared to B6 T_FH_ cells, W.Yaa T_FH_ cells exhibited a distinctive gene expression signature dominated by the cytokine, cell cycle, lysosome and mitochondrial pathways. Importantly, many orthologs of these genes were also differentially expressed in the T_FH_ cells from SLE patients compared to healthy controls. 2DG treatment reversed these distinctive pathways in W.Yaa T_FH_ cells to the B6 levels, including the expression of orthologs that are overexpressed in the T_FH_ cells from SLE patients. These common genes control the cell cycle, cytokine signaling, iron hemostasis, as well as galectins, all of which are critically involved in T_FH_ cell differentiation and functions. These results thus suggest that the inhibition of glycolysis reprograms the expression of disease-relevant genes in lupus T_FH_ cells that regulate their enhanced function. Notably, this conclusion was supported by adoptive transfers of T_FH_ cells from 2DG-treated mice, which delayed renal pathology, autoantibody production and T_FH_ cell expansion in two different lupus models.

We also showed that 2DG increased DNA methylation in W.Yaa T_FH_ cells, including in critical genes differentially expressed between W.Yaa and B6 T_FH_ cells, such as *Blimp1* and *Ubash3a*, which control T_FH_ cell differentiation and TCR/IL-2 signaling, respectively, as well as cytokine genes *Ifng, Il10* and *Lag3*, which control T_FH_ function. This suggests that glycolysis at least partially regulates the T_FH_ epigenome. Chromatin accessibility controls T_FH_ cell differentiation^63^, and we have reported an increased histone methylation in lupus T_FH_ cells^5^. DNA methylation regulates gene expression in the glycolytic pathway^64^. Further, lysine acetyltransferase KAT6A mediates an increased expression of glycolytic genes in CD4^+^ T cells of lupus-prone MRL/*lpr* mice, in which treatment with a KAT6A inhibitor reduced disease activity^65^. The present study is the first to show that the inhibition of glycolysis increases DNA methylation in T cells, which was associated with the differential expression of genes regulating T_FH_ cell differentiation and function. The mechanism by which glycolysis regulates DNA methylation is unknown. However, our results showed a critical role of the inhibition of glycolysis in restoring mitochondrial homeostasis in lupus T_FH_ cells, which in turn may promote DNA methylation, whereby reduced mitochondrial ATP production alters DNA methylation^66^.

Reprogramming of mitochondrial function by 2DG in W.Yaa T_FH_ cells was documented by both the restored expression of mitochondrial and respiratory genes and increased mitochondrial metabolites (e.g. carnitine, succinate). Defective mitochondria are a major driver of T cell dysfunction in SLE^67^, characterized by impaired mitophagy^68^. We have also reported a transcriptional signature associated with defective mitochondrial functions, including mitophagy, in TC T_FH_ cells^5^, suggesting that it may be a feature shared by lupus T_FH_ cells. We also showed that W.Yaa T_FH_ cells presented a lysosomal / autophagy gene signature that was restored by the inhibition of glycolysis, most likely independently from the mTOR pathway. Several studies have shown that dysfunctional mitochondria alter the lysosome ^69^. In T cells in particular, mitochondrial defects critically impair the lysosome and promote inflammatory phenotypes that can be reversed by restoring NAD^+^ levels^70^. Accordingly, W.Yaa T_FH_ cells showed reduced mROS and NAD^+^ levels, and our results suggest that low levels of complex I might be mainly responsible for the lower production of these two critical metabolites. mROS is required for autophagy flux^71^ and NAD^+^ acts bidirectionally as a regulator and target of autophagy^72^. Canonical markers of macroautophagy remained unchanged in W.Yaa T_FH_ cells or by the inhibition of glycolysis. Instead, we found increased LAMP2A-driven CMA in W.Yaa T_FH_ cells, which was normalized by 2DG. Furthermore, we provided evidence that TLR7 activation promoted CMA in T_FH_ cells. CMA is increased in B cells from MRL/*lpr* mice and a treatment with the P140 peptide to reduce CMA activity ameliorated clinical manifestations in this model as well as in patients with SLE^73^. It has been proposed that the inhibition of CMA in B cells reduces the autoantigen load on MHCII molecules, consequently lowering the activation of autoreactive T cells. More directly related to the present study, CMA promotes T cell activation by selectively deleting negative regulators of TCR signaling^74^. Thus, increased CMA activity in W.Yaa T_FH_ cells is thought to enhance TCR activation, consistent with the transcriptomic signature of lupus T_FH_ cells^5^ (and this study). Furthermore, our results support a model in which a defective lysosomal function in W.Yaa T_FH_ cells promotes glycolysis by increasing the expression of the rate-limiting HK2. By blocking HK2 activity, 2DG rewired mitochondrial homeostasis, restored lysosomal functions and reduced CMA, thereby reducing T_FH_ cell responses, which in turn improved the health status of lupus-prone animals. Overall, these preclinical results provide a mechanistic basis by which uncontrolled glycolysis promotes T_FH_ cell expansion and increases their function in lupus-prone mice. The overlap in transcriptional signatures suggests that this model may also be relevant to human lupus T_FH_ cells. Our results also offer a strong rationale for the therapeutic potential of targeting glycolysis to limit the progression of lupus and subsequently improve patient health, at least in part by normalizing T_FH_ cell function.

## ACKNOWLEDGMENTS

We thank the staff from the University of Florida Lupus Clinic, the UF SECIM metabolomic core for technical support and the UF Molecular Pathology Core for histology services, as well as the staff of the UTHSA Flow Cytometry Shared Resource.

## AUTHOR CONTRIBUTIONS

Conceptualization, A.E., M.M., and L.M. Methodology, A.E., Y.G. and S.C.C, Investigation, A.E., Y.G., S.C.C, Y.P.P., L.P., Z.Y., and W.L.C; Writing – Original Draft, A.E., and L.M. Writing – Review & Editing, Y.G., S.C.C., M.M. and L.M.; Resources, E.S.S.; Funding Acquisition, M.M., and L.M.; Supervision, M.M. and L.M.

## STAR methods

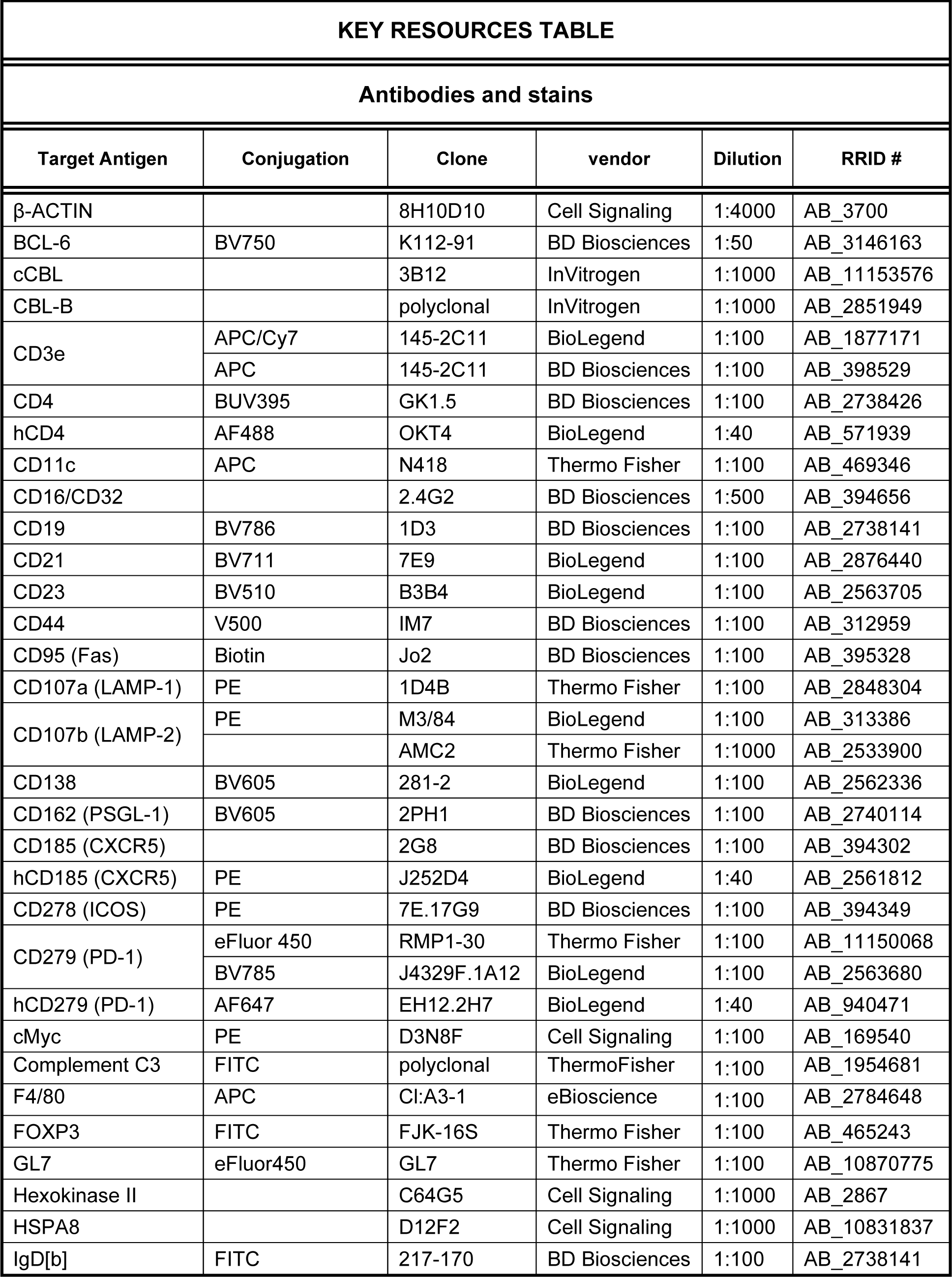
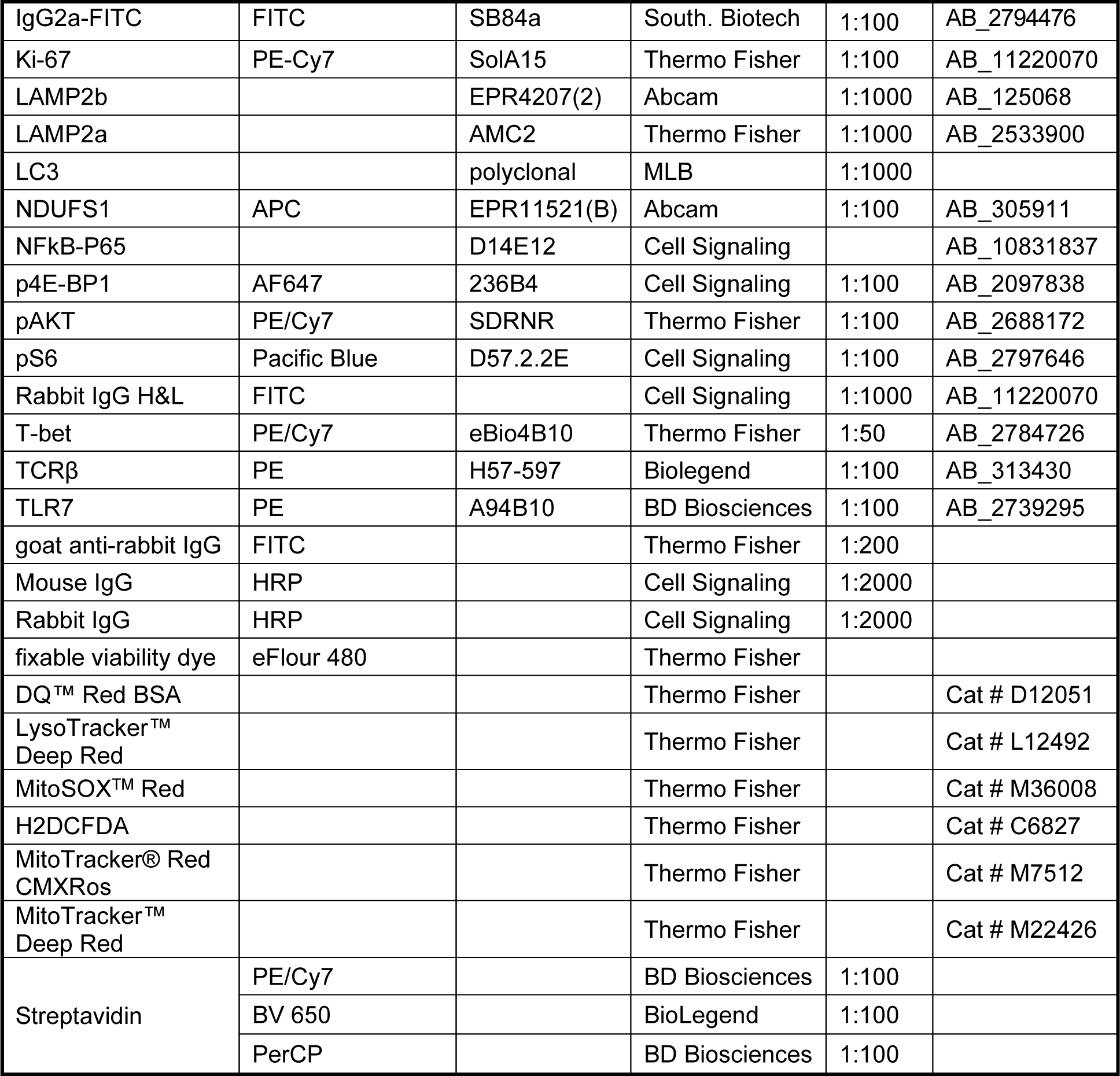
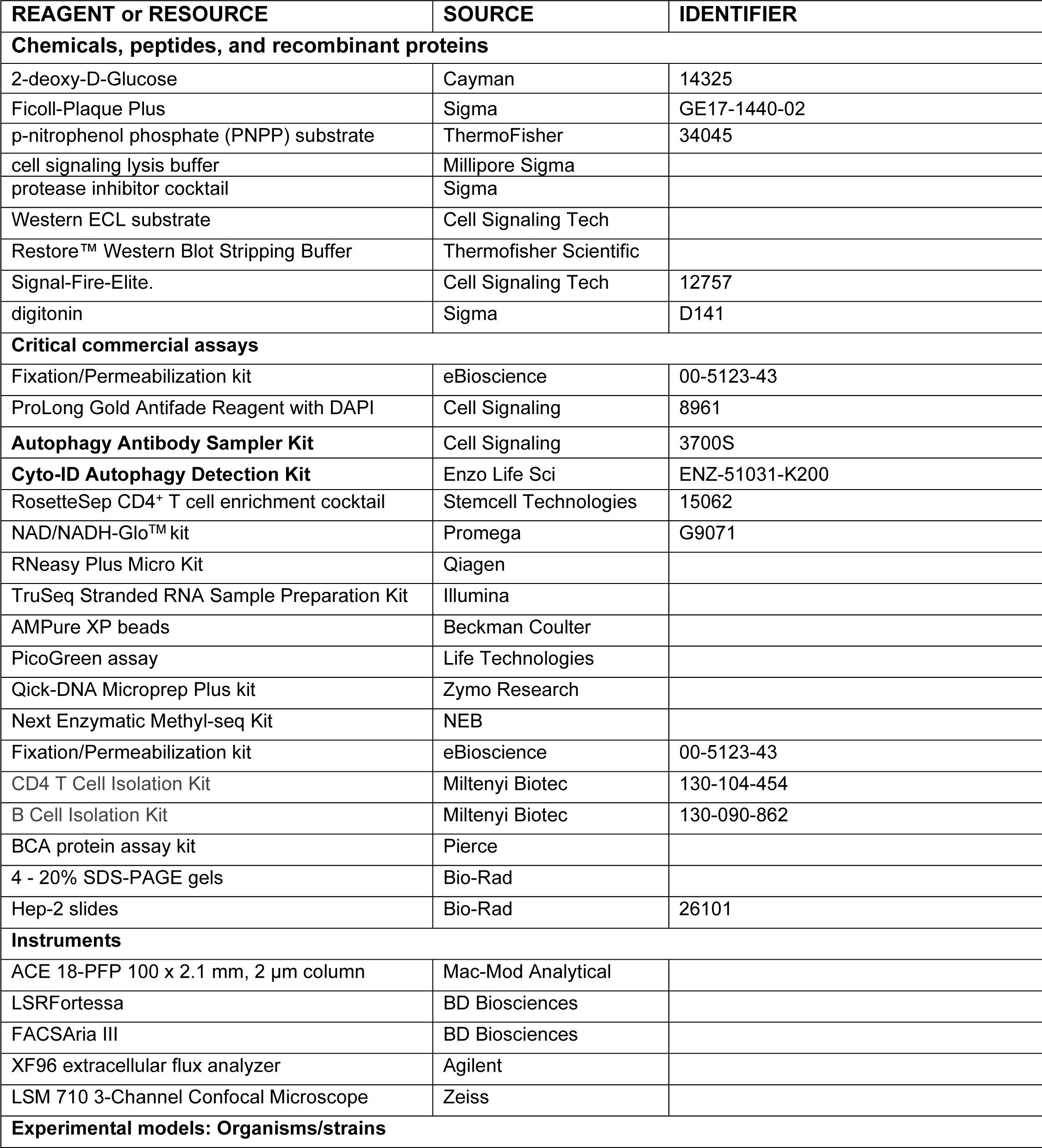
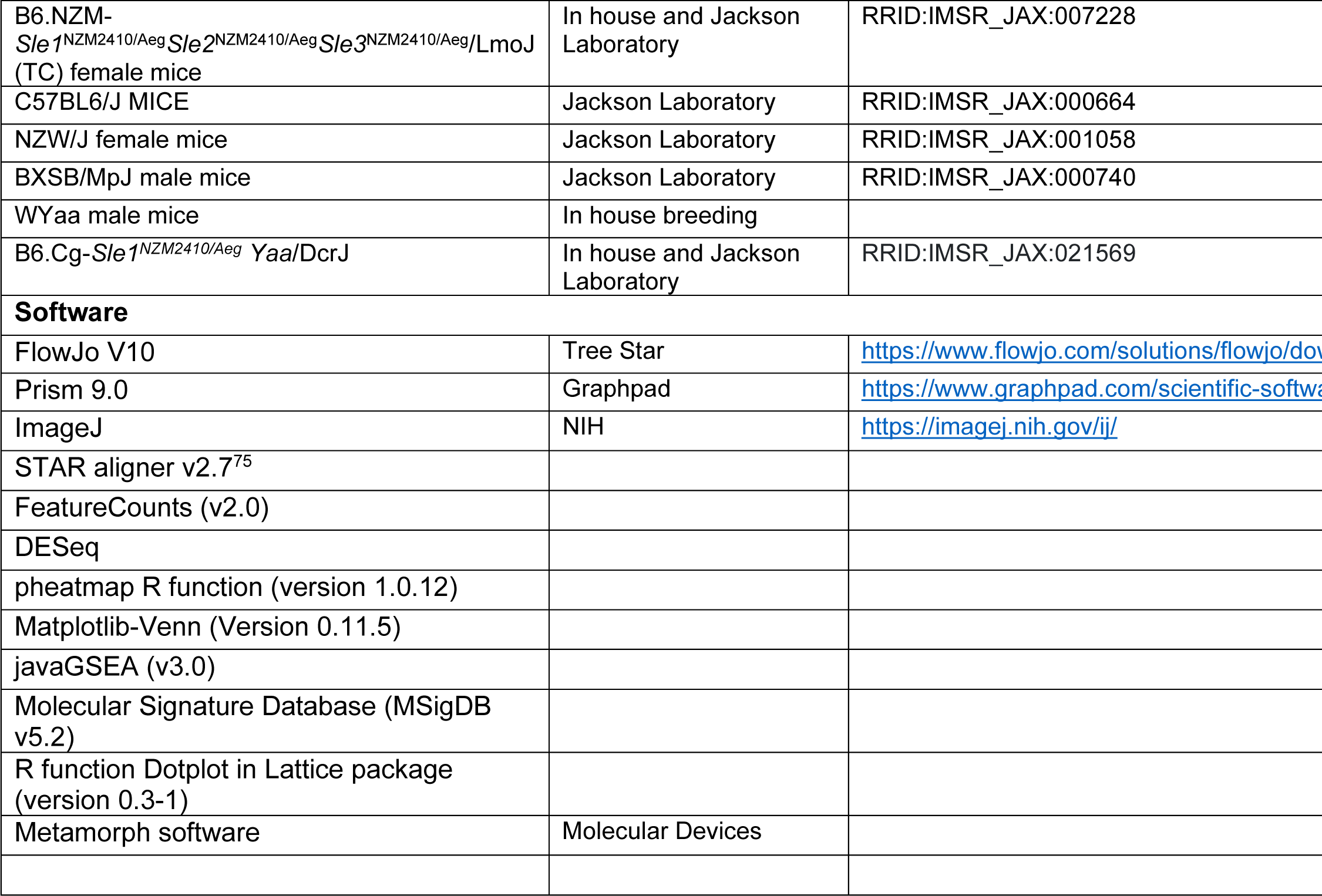

## LEAD CONTACT

Further information and requests for resources and reagents should be directed to and will be fulfilled by the lead contact, Dr. Laurence Morel.

## MATERIALS AVAILABILITY

RNASeq raw datasets are available under NCBI BioProject accession numbers PRJNA1089578 (mouse) and PRJNA1089580 (human). All other data reported in this paper will be shared by the lead contact upon request.

## EXPERIMENTAL MODEL

NZW/J females and BXSB/MpJ males were bred to produce F1 male progeny (W.Yaa). The B6.NZM-*Sle1*^NZM2410/Aeg^*Sle2*^NZM2410/Aeg^*Sle3*^NZM2410/Aeg^/LmoJ (TC), B6.*Sle1* and B6.*Sle1*.Yaa congenic mice have been previously described ^51,76^. W.Yaa mice and their controls were males, and all TC mice and their controls were females. Cohorts of 10 - 12 week-old W.Yaa mice or 18 – 20 week old TC mice, corresponding to an early stage of disease for each strain with anti-dsDNA IgG production without renal pathology, and age-matched control mice were orally treated or not with 2DG (6 mg/ml) in drinking water. Mice were euthanized after 4 weeks of treatment or earlier if proteinuria reached > 300 mg/dl for 2 consecutive weeks. For T_FH_ cell transfers, 1.5 x 10^6^ T_FH_ cells from individual untreated mice or from pooled 2DG-treated mice were injected in the retro-orbital sinus of either 6-week-old W.Yaa or 2 month-old TC mice (matched strain between donor and recipients). The autoimmune phenotypes of recipients were evaluated at 14 weeks of age for W.Yaa mice and 7 month-old TC mice. The difference in age between the two strains corresponds to the slower disease progression in TC mice.

Mice were maintained in SPF conditions at the University of Florida (UF) or at the University of Texas Health San Antonio (UTHSA). This study was carried out in accordance with the guidelines from the Guide for the Care and Use of Laboratory Animals of the Animal Welfare Act and the NIH (National Research Council) Guide for the Care and Use of Laboratory Animals. All animal protocols were approved by the Institutional Animal Care and Use Committee of the University of Florida and the University of Texas Health San Antonio (IACUC 202009466 and 20220032AR, respectively).

## HUMAN SUBJECTS

PBMCs were collected from SLE patients and healthy controls (HC) at the University of Florida Lupus Clinic according to an approved protocol (UF IRB 201300225). The recruitment and study were carried out in accordance with the World Medical Association Declaration of Helsinki: Ethical Principles for Medical Research Involving Human Subjects. Written informed consent was obtained from all study participants, all of whom were all females, and their demographics are listed in Supplementary Table S1.

## METHODS DETAILS

### ANTIBODY MEASUREMENT

Antinuclear antibodies (ANAs) were detected on Hep-2 cell-coated slides in sera diluted 1:40. Anti-RNA IgG were detected by ELISA in sera diluted 1:100^77^.

### FLOW CYTOMETRY AND CELL SORTING

Flow cytometry was performed on splenocyte suspensions prepared as previously described^12^. Human PBMCs were isolated from whole blood using Ficoll-Plaque Plus. Cells were stained with fluorochrome-conjugated antibodies. Dead cells were excluded with fixable viability dye. For intracellular staining, cells were fixed and permeabilized using the FOXP3/Transcription Factor Staining Buffer. Data were acquired using an LSRFortessa or FACSymphony A5, and analyzed with FlowJo V10 (BD Bioscience). Gating strategies for CD4^+^ T cell and B cell subsets are shown in Figure S10A for mouse and Figure S11 for human cells. Murine T_FH_ cells were gated as FOXP3^-^ CD44^+^PD-1^+^CXCR5^+^BCL6^+^ PD-1^+^PSGL1^lo^CD4^+^ T cells, T_FR_ as FOXP3^+^ PD-1^+^CXCR5^+^BCL6^+^ PD-1^+^PSGL1^lo^CD4^+^ T cells, and T_EXFH_ as FOXP3^-^CD44^+^PD-1^+^CXCR5^-^BCL6^-^PD-1^+^PSGL1^lo^CD4^+^ T cells. Cell sorting was performed with a FACSARIA III cytometer. For RNASeq and metabolomic analyses, murine T_FH_ cells were sorted as CD4^+^CD44^+^PD-1^+^PSGL-1^lo^ cells and T_N_ cells were sorted CD4^+^CD44^-^ cells. Purity was > 95% (Figure S10B). Human PD-1^+^CXCR5^+^ T_FH_ cells were sorted from CD4^+^ T cells isolated from PBMCs by negative selection with the RosetteSep CD4^+^ T cell enrichment cocktail. Purity after sorting was approximately 90% (Figure S12). The Cyto-ID Autophagy Detection Kit was used on splenocytes treated with Cyto-ID diluted 1:1000 and fixed with 10% formalin. Mean fluorescence intensity was measured by flow cytometry.

### METABOLIC MEASUREMENTS

Mitochondrial stress assays were conducted on an XF96 extracellular flux analyzer on magnetic-bead purified CD44^+^ CD4^+^ T cells, CD4^+^ T cells or B cells. Data was normalized using a coupled Biotek Cytation 1 imager or to an age-matched B6 sample in each assay. The NAD^+^/NADH ratio was measured in CD44^+^ CD4^+^ T cells with the NAD/NADH-Glo^TM^ kit. Metabolomic analyses of sorted murine (7.5 - 10 x 10^5^ cells) and human T_FH_ cells (1 - 4 x 10^5^ cells) were performed at the University of Florida SECIM metabolomic core. Global metabolomics profiling was performed on a Thermo Q-Exactive Oribtrap mass spectrometer with Dionex UHPLC. All samples were analyzed in positive and negative heated electrospray ionization with a mass resolution of 35,000 at m/z 200 as separate injections. Separation was achieved on an ACE 18-pfp 100 x 2.1 mm, 2 µm column with mobile phase A as 0.1% formic acid in water and mobile phase B as acetonitrile. The flow rate was 350 µl/min with a column temperature of 25°C. 4 µl were injected for negative ions and 2 µL for positive ions. Metabolites were identified by comparison to the library of purified standards or putatively annotated by their enrichment in significant pathways and by the m/z match with an accuracy of 10 ppm. These metabolites were labeled with asterisks (*) in heatmaps.

### RNA-SEQ ANALYSIS

RNA was isolated from FACS-sorted T_FH_ and T_N_ cells using the RNeasy Plus Micro Kit. Raw sequencing reads were mapped to the reference mouse (GRCm38) or human (GRCh38.83) genomes using STAR v2.7.5c. DESeq2 was used to identify the differentially expressed genes (DEGs) based on the criteria (TPM > 1, FDR < 0.05, fold change > 1.5). Gene set enrichment analysis was performed using DAVID (https://david.ncifcrf.gov).

### GENOME-WIDE DNA METHYLATION ANALYSIS

Genomic DNA was isolated from FACS-sorted T_FH_ cells using the Qick-DNA Microprep Plus kit. Bisulfite-converted DNA libraries were constructed using the Next Enzymatic Methyl-seq Kit and sequenced on an Illumina NovaSeq 6000 platform. Raw sequence reads were first trimmed to remove poor quality nucleotides and the high-quality reads were then mapped to the mouse reference genome (GRCm38) using Bismark v0.22.3, yielding at least 17-fold whole genome coverage. Differentially methylated regions (DMRs) were identified using the package DSS and genomic annotation of the DMRs was performed using BEDTools v2.29.2 ^78^.

### Western blotting

Total protein extracted from magnetic bead-purified CD4^+^CD62L^+^ and CD4^+^CD44^+^CD62L^-^ T cells was quantified using the BCA method. Immunoblotting was performed according to a standard protocol with primary antibodies and HRP-conjugated anti-rabbit or anti-mouse IgG secondary antibodies. Signal was detected using Signal-Fire-Elite. Image J software was used for densitometric analysis. For repeated measurements of different proteins on the same membrane, PVDF stripping buffer was applied and the membrane was re-blocked before re-probing.

### CONFOCAL MICROSCOPY

Sorted CD4^+^CD44^+^CD62L^-^ T cells (10^6^ cells/ well) were fixed with 4% paraformaldehyde, permeabilized with digitonin, then stained with primary antibodies against LAMP2a and LC3, and PE-conjugated anti-mouse LAMP-1. After washing, cells were incubated with FITC-conjugated goat anti-rabbit IgG. Stained cells transferred to cytoslides were incubated with ProLong Gold Antifade Reagent with DAPI and visualized by a Confocal Microscope equipped with a 60X oil immersion objective. A scan resolution of 2048 x 2048 was used to image acquisition.

### RENAL PATHOLOGY

Paraformaldehyde-fixed paraffin kidney sections were stained with periodic acid Schiff (PAS). The type and extent of renal lesions were evaluated using a modification of the International Society of Nephrology and Renal Pathology Society classification of lupus nephritis and the NIH activity and chronicity indices in a blinded manner by a pathologist (WLC). Renal parenchymal components, including glomeruli, vessels, tubules, and interstitium, distributed throughout the renal sections were assessed. At least 100 glomeruli per kidney were examined for the presence of mesangial expansion, mesangial and endocapillary hypercellularity, crescents, and glomerulosclerosis. Glomerulonephritis scores were calculated as the sum of the 6 criteria divided by 100. Acute tubular injury lesions, including tubular dilatation, epithelial attenuation, and tubular casts were quantified on a scale of 1 to 4. Glomerular area was calculated with the Metamorph software. IgG2a and C3 immune complexes as well as T cell and macrophage infiltration were detected in frozen sections as previously described^15^.

### STATISTICAL ANALYSIS

Statistical analyses were performed using GraphPad Prism 9.0 software. Unless otherwise stated, differences between groups were evaluated by ANOVA with correction for multiple testing or unpaired or paired *t* tests, as indicated in the figure legends. The corresponding nonparametric tests were used when the data distribution deviated from normality. Results were expressed as means ± standard errors of the mean (SEM). Statistical significance levels were set at *: *P* < 0.05, **: *P* < 0.01, ***: *P* < 0.001 and ****: *P* < 0.0001.

