## supplemental table and figures for "Glycolysis inhibition functionally reprograms T follicular helper cells and reverses lupus": bioRxiv supplemental.pdf

**Supplementary Table S1.** Demographics of the human subjects used in this study (all females).

| Ethnicity | Age | | |
|---|---|---|---|
| **SLE patients** | | | |
| | | **Drugs (mg / d)** | **SLEDAI** |
| W | 44 | CellCept 250 | 0 |
| B | 58 | CellCept 1500 | 4 |
| W/H | 68 | Prednisone (15) | 9 |
| W | 45 | CellCept 1000 | 2 |
| W | 43 | Imuran | 2 |
| B | 51 | Imuran | 5 |
| B | 38 | Tacrolimus | 18 |
| W | 42 | CellCept 1000 | 0 |
| B/H | 25 | CellCept 1000 Prednisone (40) | 8 |
| W | 45 | CellCept 1000 Prednisone (40) | 8 |
| W | 44 | CellCept 250 | 0 |
| **Healthy controls** | | | |
| B | 26 | | |
| W | 31 | | |
| B | 42 | | |
| W | 23 | | |
| B | 26 | | |
| W | 21 | | |
| B | 26 | | |
| W/H | 25 | | |
| W | 25 | | |
| W | 29 | | |

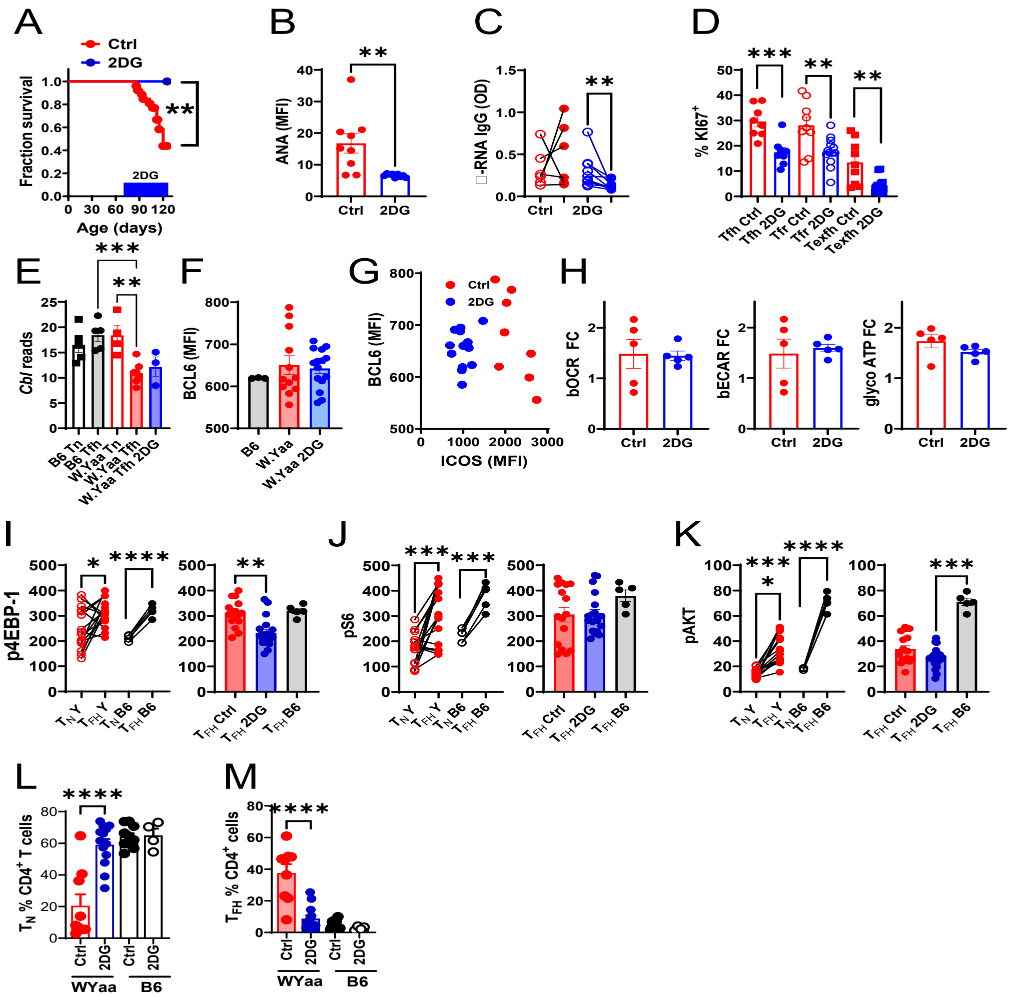


**Figure S1**. Immunopathology and cellular effects of 2DG in W.Yaa mice. (**A**) Survival of 2DG-treated (N = 10) and control (N = 16) W.Yaa mice compared with a Kaplan-Meyer test. The blue box indicates the 2DG treatment. (**B**) Terminal serum ANA (N = 9, *t* test). (**C**) Serum anti-RNA IgG before and after treatment (N = 6 -10, paired *t* test). (**D**) Frequency of proliferating KI67$^{+}$ $T_{FH}$, $T_{FR}$ and $T_{EXFH}$ cells from control and 2DG treated W.Yaa mice (N = 9 – 13, Dunnett's T3 multiple comparisons tests). (**E**) *Cbl* expression in $T_N$ and $T_{FH}$ cells from B6 and W.Yaa mice (RNAseq results, N = 3 – 7). (**F**) BCL-6 expression and (**G**) correlation with ICOS expression in $T_{FH}$ cells (MFI measured by flow cytometry) in the indicated strains. (**H**) Mitochondrial stress assay in B cells from 2DG-treated or control W.Yaa mice, with from left to right basal OCR, basal ECAR, and glycolytic ATP calculated as fold-change from an untreated B6 control (N = 5). (**I** – **K**) mTOR activation in W.Yaa $T_{FH}$ cells. Graphs on the left show untreated W.Yaa (Y) and B6 $T_N$ and $T_{FH}$ cells from the same mice compared with paired *t* or Wilcoxon matched-pair signed rank tests. Graphs on the right compare the same parameters in $T_{FH}$ cells from untreated controls or treated with 2DG (N= 6 – 19) W.Yaa mice, and age-matched untreated B6 (N = 5). (**I**) p4EBP1; (**J**) pS6; (**K**) pAKT ser 473 all shown as MFI and compared with Dunnett's T3 multiple comparisons tests. Frequency of $T_N$ (**L**) and CD4$^{+}$CD44$^{+}$PD-1$^{+}$PSGL1$^{lo}$ $T_{FH}$ (**M**) cells from W.Yaa (N = 9 – 13)and B6 (N = 4 -10) mice treated or not with 2DG that were used in OMICs experiments, compared with Dunnett's T3 multiple comparisons tests. Mean + SEM, *: P < 0.05, **: P < 0.01; ***: P < 0.001; ****: P < 0.0001.

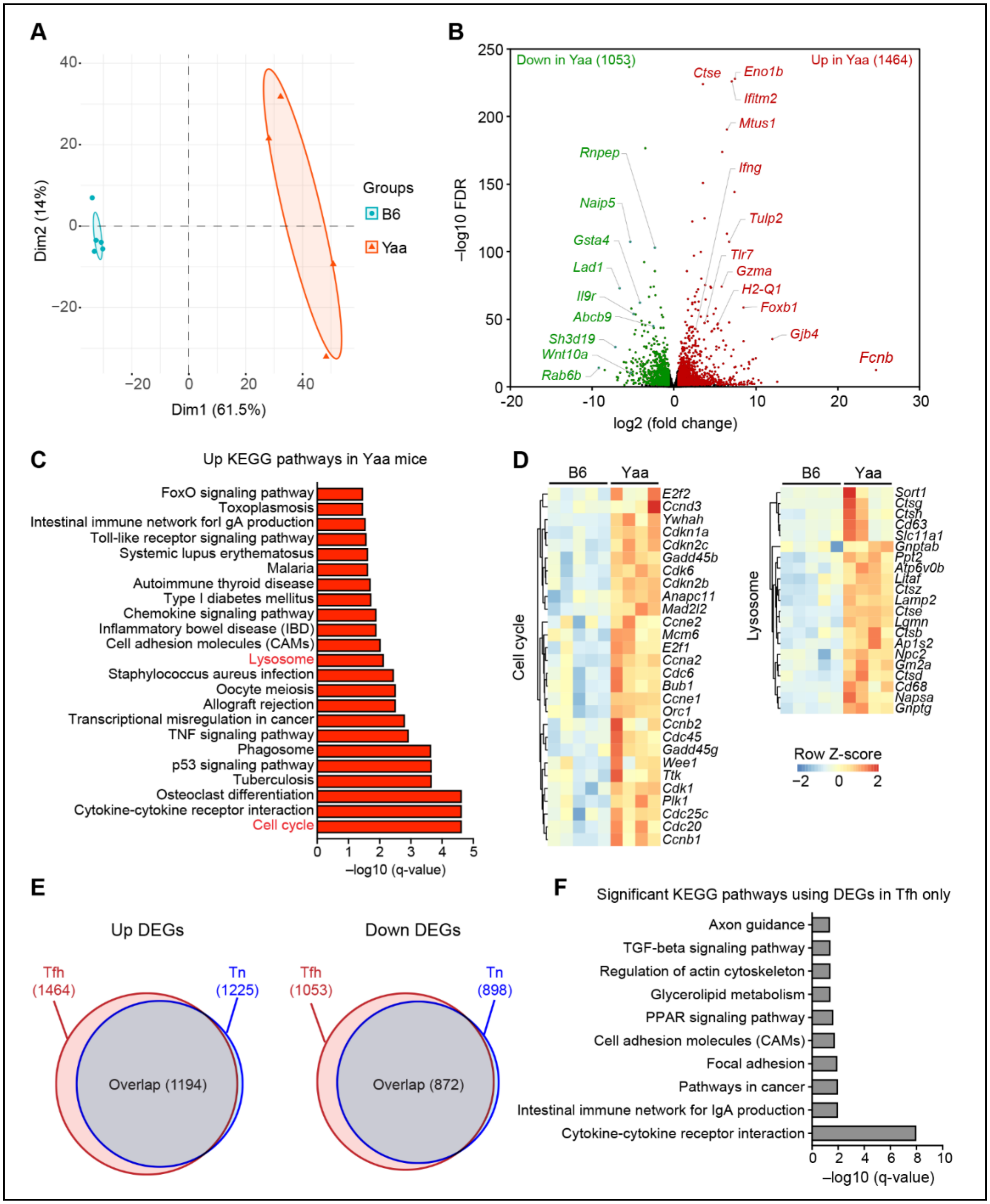


**Figure S2.** W.Yaa Tfh cells presented a distinct gene expression profile from B6 Tfh cells. RNASeq was performed on $T_N$ and $T_{FH}$ cells sorted from 14 weeks old W.Yaa and B6 mice (N = 4 -5 per group). (**A**) PCA analysis showing the W.Yaa and B6 $T_{FH}$ cell clusters. (**B**) Volcano plots of DEGs between W.Yaa and B6 $T_{FH}$ cells. Arrows point to *Trl7,* a gene duplicated in W.Yaa mice, and *Eno1b* encoding for a glycolytic enzyme. (**C**) Upregulated KEGG pathways in W.Yaa $T_{FH}$ cells, and heatmaps of the Cell cycle and Lysosome pathways (**D**). (**E**) The majority of DEGs between W.Yaa (red) and B6 (bleue) $T_{FH}$ cells also differed between the corresponding $T_N$ cells. **F**. KEGG pathways uniquely differently expressed in $T_{FH}$ cells between the two strains.

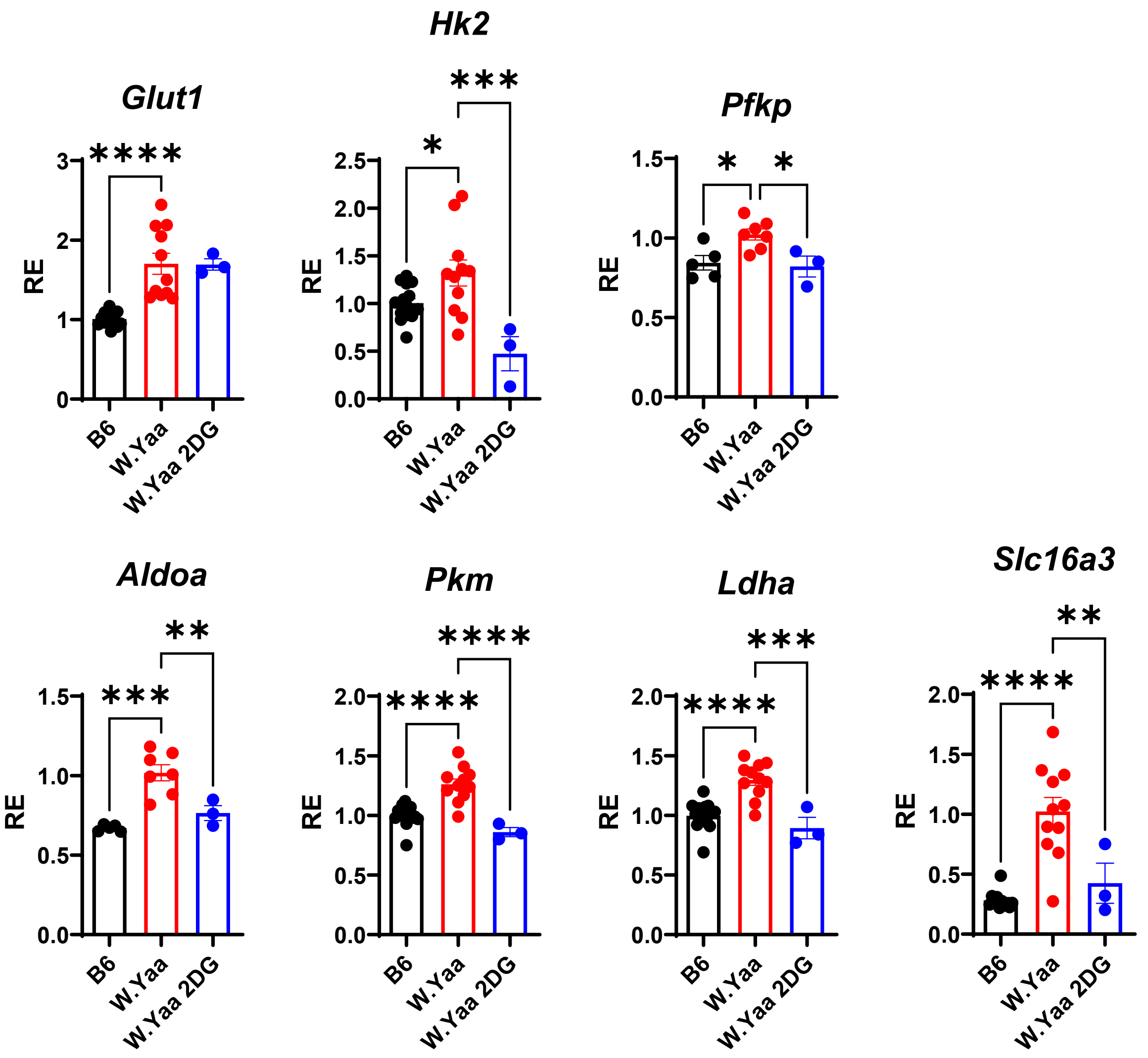


**Figure S3.** W.Yaa $T_{FH}$ cells express higher levels of genes in the glycolytic pathway that are downregulated in mice treated with 2DG. Normalized RNASed reads with mens + SEM compared with Šídák's multiple comparisons tests. *: P < 0.05, **: P < 0.01; ***: P < 0.001; ****: P < 0.0001. N = 3 – 15.

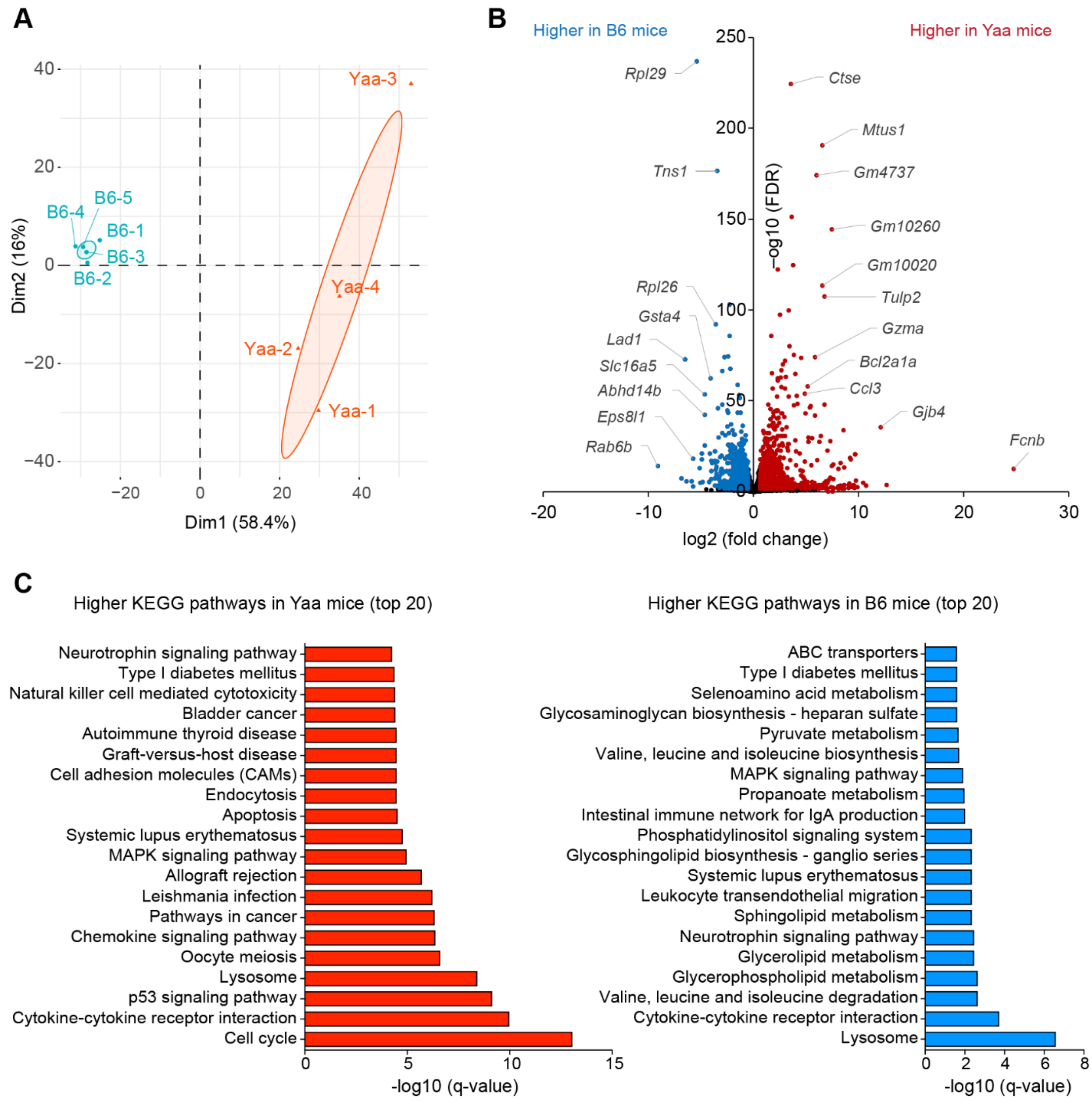


**Figure S4.** W.Yaa and B6T$_N$ cells presented a distinct gene expression profile. (**A**) PCA analysis showing the W.Yaa and B6 T$_N$ cell clusters. (**B**) Volcano plots of DEGs between W.Yaa and B6 T$_N$ cells. (**C**) Upregulated KEGG pathways in W.Yaa T$_N$ cells (left) and B6 T$_N$ cells (right).

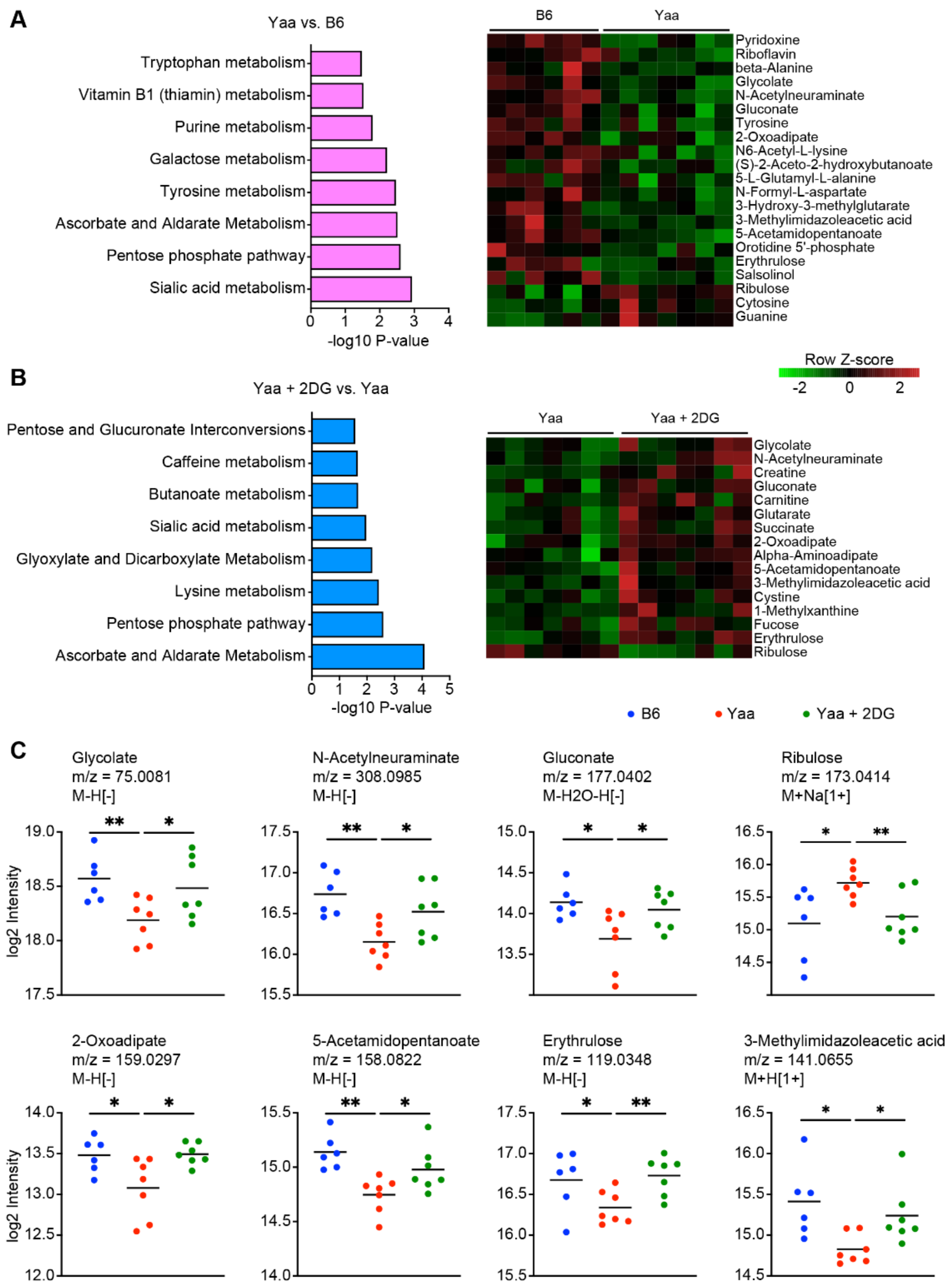


**Figure S5.** The inhibition of glycolysis normalized the W.Yaa Tfh metabolome. Untargeted metabolomics was performed on $T_{FH}$ cells from B6 (N = 6) as well as 2DG-treated and control W.Yaa (N = 7 each) mice**.** Pathway analysis and heatmap of metabolites with significantly differing intensity between $T_{FH}$ cells from B6 and W.Yaa control mice (**A**) and from 2DG-treated and control W.Yaa mice (**B**). (**C**) Selected metabolites in the three groups, with their respective m/z values and adduct ions. Horizontal bars indicate means. Comparisons were made with Dunnett's T3 multiple comparisons tests to the W.Yaa control samples. *: P < 0.05; **: P < 0.01.

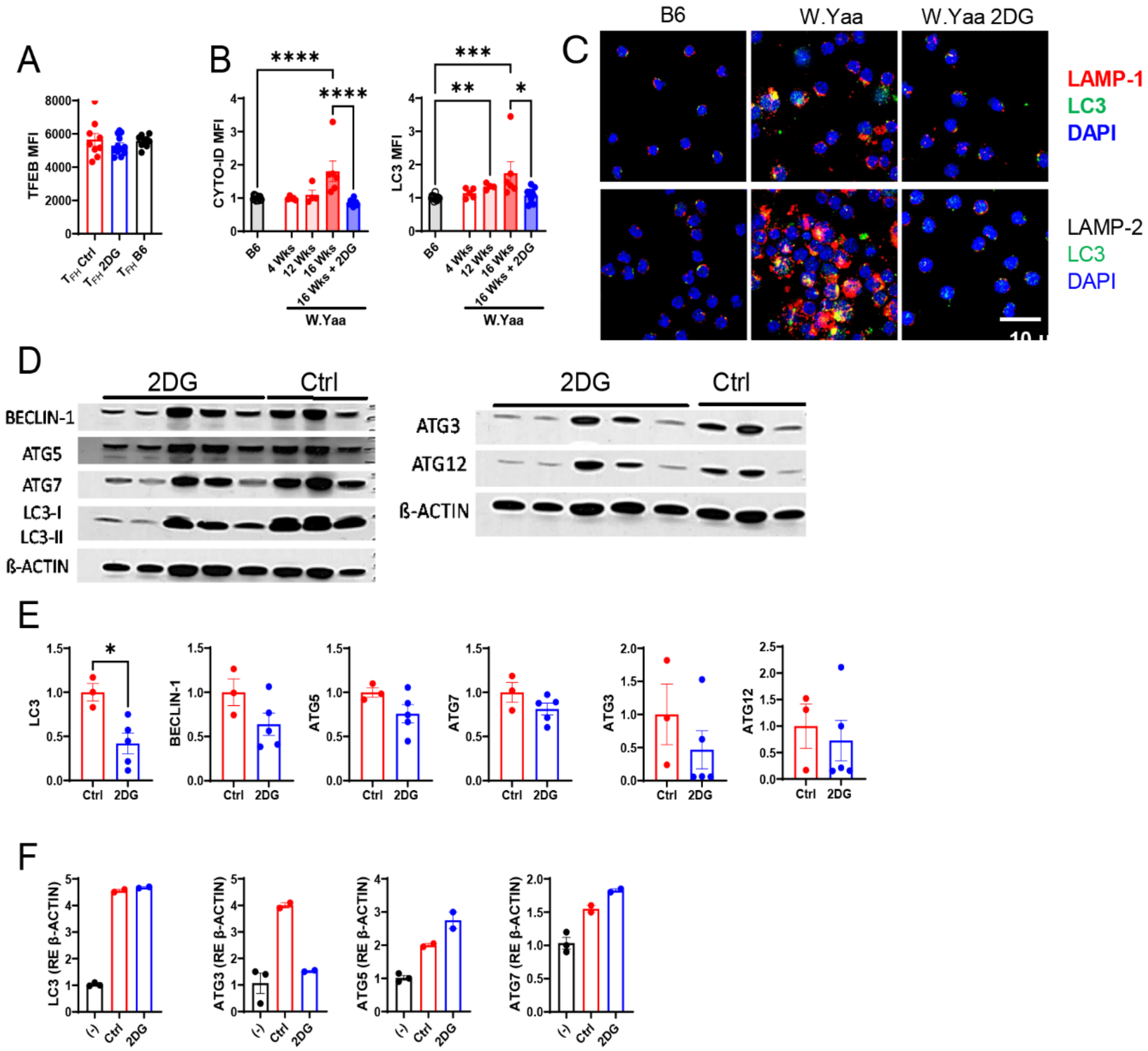


**Figure S6.** Autophagy analysis in W.Yaa $T_{FH}$ cells. (**A**) TFEB expression in $T_{FH}$ cells from W.Yaa mice treated with 2DG or controls, or age-matched untreated B6 (N = 9 - 12). (**B**) CYTO-ID and LC3 expression in Tfh cells from 4, 12 and 16-weeks old W.Yaa and B6 mice, as well as from 16 weeks old W.Yaa mice after 4 weeks of 2DG treatment. The B6 values were similar between age groups and were pooled together. Values shown as MFI measured by flow cytometry compared with Dunnett's T3 multiple comparisons tests. (**C**) Representative confocal images of defective autophagolysosome in W.Yaa $CD4^+$ T cells. Top: W.Yaa $CD4^+$ T cells accumulated LC3 unfused with LAMP1-stained lysosome. Bottom: W.Yaa $CD4^+$ T cells presented large LC3-LAMP2 aggregates. 2DG normalized the autophagolysosome of W.Yaa $CD4^+$ T cells to a B6-like phenotype. (**D**) Western blot analysis of $CD44^+CD4^+$ T cells from 2DG-treated and control W.Yaa mice probed for BECLIN-1, ATG5, ATG7, LC3, ATG3 and ATG12, with quantification relative to β-ACTIN shown in (**E**) compared with *t* tests. (**F**) B6 $CD4^+$ T cells were activated with anti-CD3 and CD28 for 24 h with control medium or 2DG and probed by Western blot for LC3, ATG3, ATG5 and ATG7. (-) indicate non-activated cells. N = 2 - 3. Mean + SEM, *: $P < 0.05$; **: $P < 0.01$; ***: $P < 0.001$; ****: $P < 0.0001$.

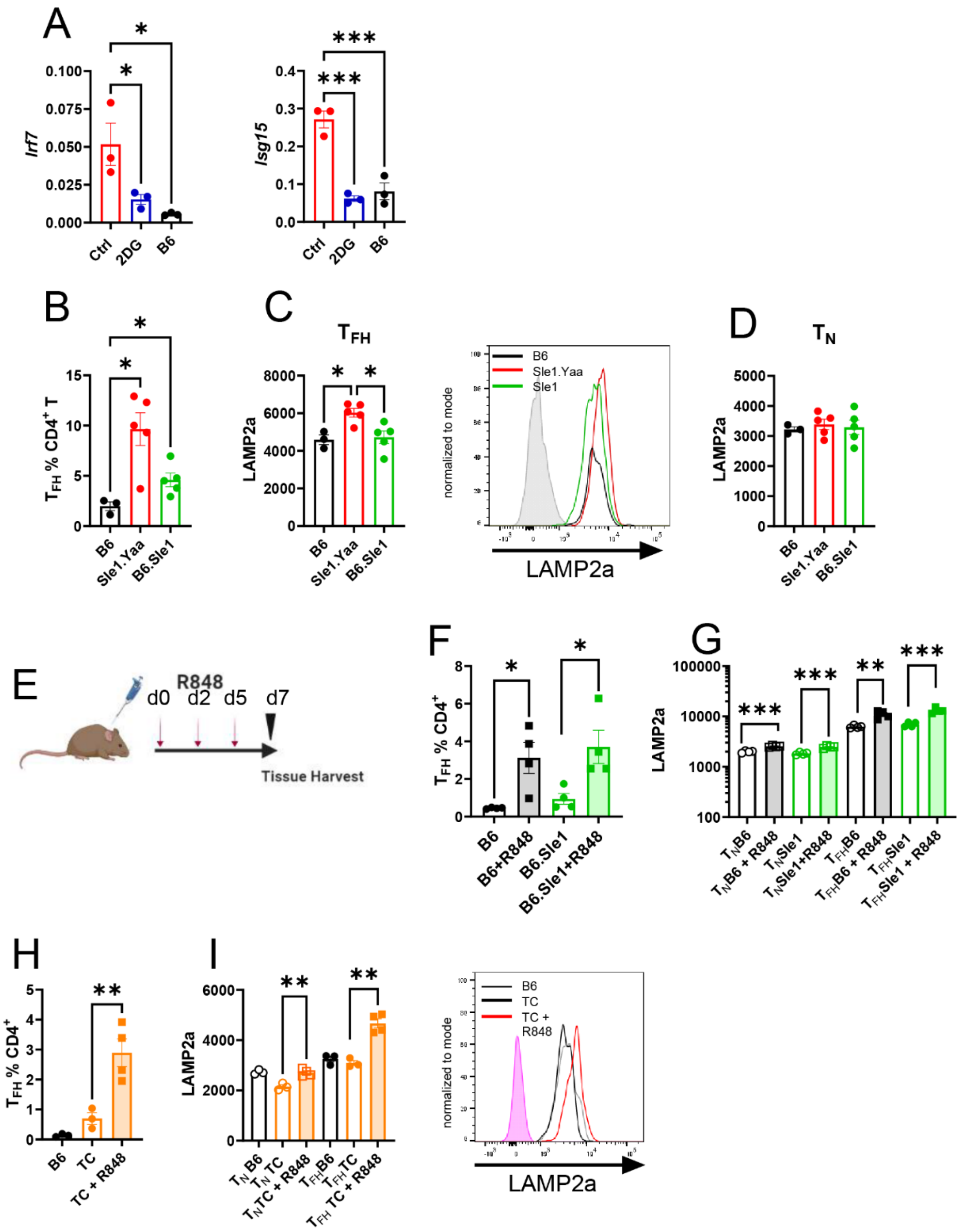


**Figure S7.** TLR7 activation increased LAMP2a expression in $T_{FH}$ cells from lupus-prone mice. (**A**) 2DG reduced the expression of interferon-inducible genes *Irf7* and *Isg15* in W.Yaa CD4$^{+}$ T cells. (**B** - **D**) LAMP2a expression in B6, B6.*Sle1*.Yaa and B6.*Sle1* CD4$^{+}$ T cells: Frequency of $T_{FH}$ cells (**B**) and LAMP2a expression in these cells (**C**) with a representative overlay histograms on the right. (**D**) LAMP2a expression in the corresponding Tn cells. (**E**) R848 treatment in B6 and B6.*Sle1* mice (**F** – **G**), and B6 and TC mice (**H** – **I**). Frequency of $T_{FH}$ cells (**F** and **H**) and corresponding LAMP2a expression (**G** and **I**). (**I**) Representative overlay histograms in $T_{FH}$ cells. Dunnett's T3 multiple comparisons tests (**A** – **D**) and *t* tests (**F** – **I**). Mean + SEM, *: P < 0.5; **: P < 0.01; ***: P < 0.001.

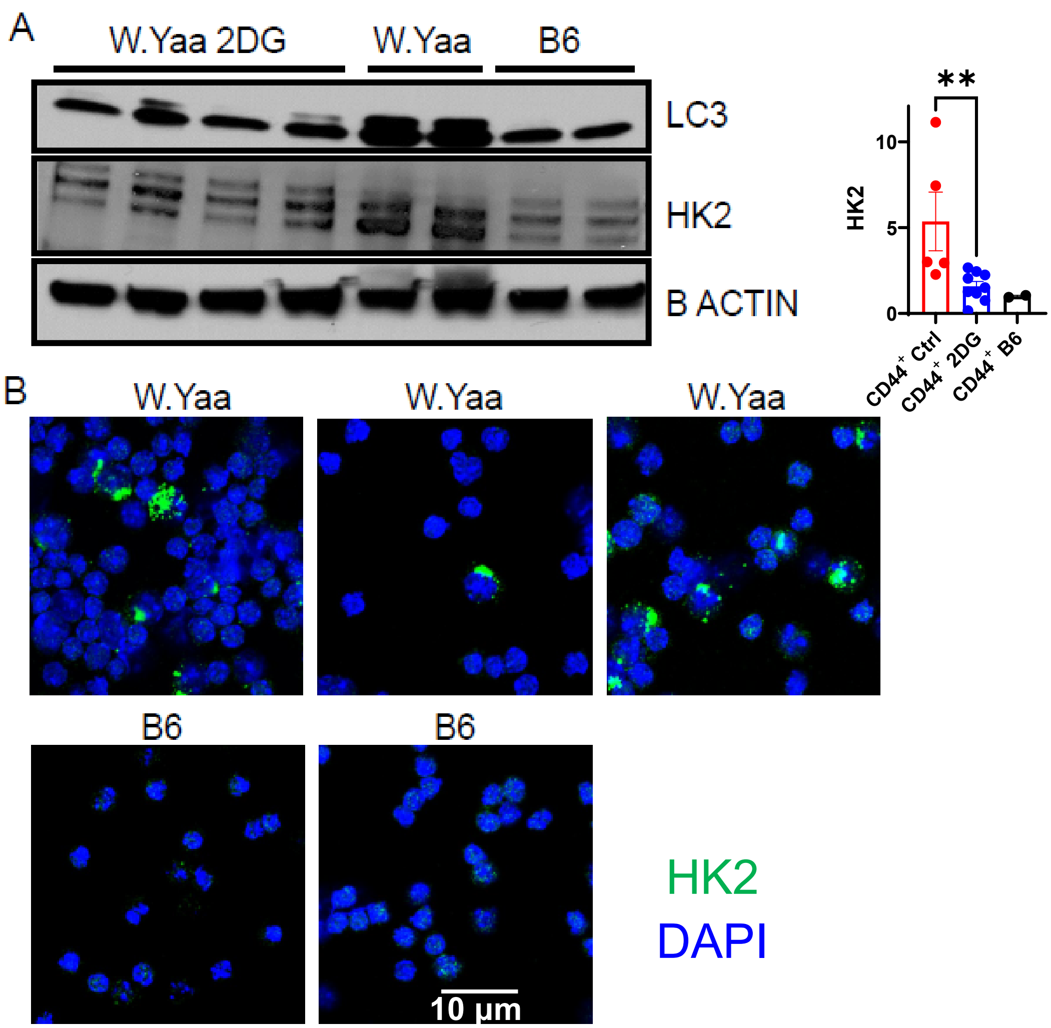


**Figure S8**. H2K expression is higher in W.Yaa than in B6 CD4$^+$ T cells and it is reversed by 2DG treatment. (**A**) Representative Western blot of LC3 and H2K, normalized to b-ACTIN in CD4$^+$ T cells from W.Yaa mice treated with 2DG and controls (N = 5 and 8, respectively), as well as from B6 age-matched controls (N = 2). HK2 quantitation on the right, compared with a Mann-Whitney test, **: $P < 0.01$. (**B**) Representative confocal images of W.Yaa and B6 CD4$^+$ T cells stained with H2K and DAPI.

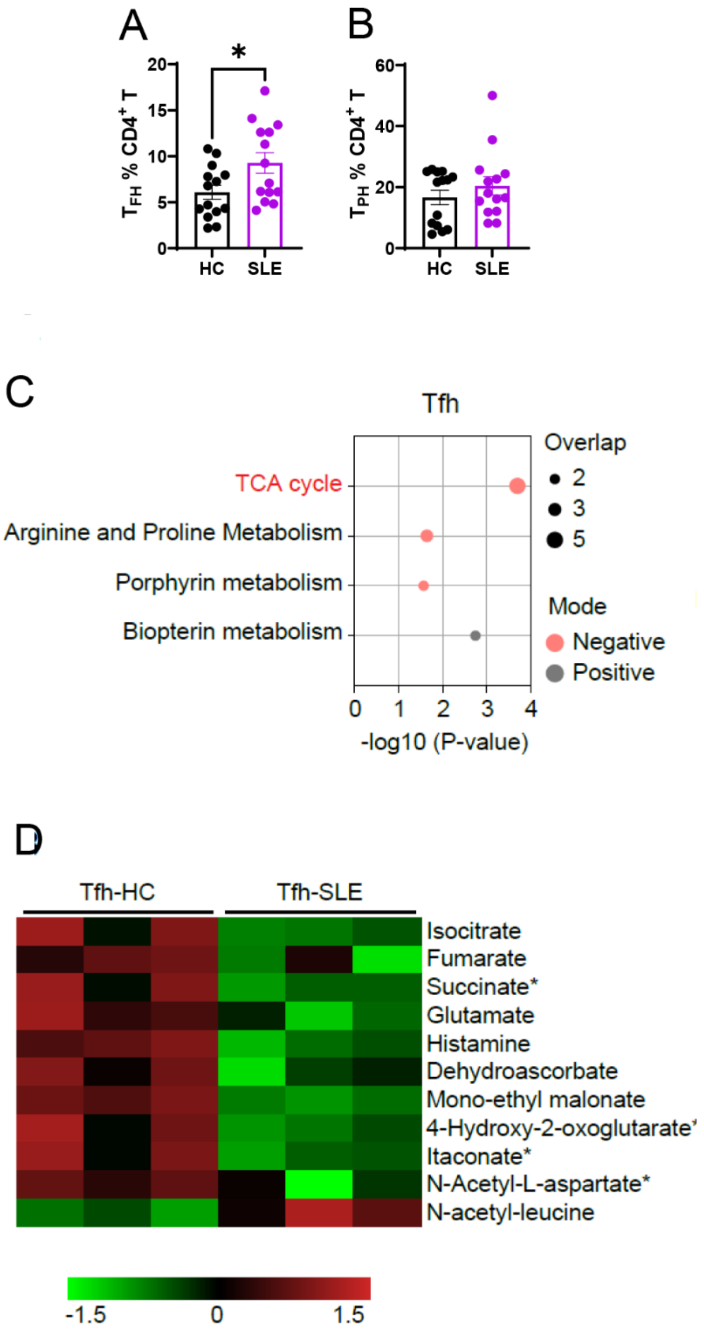


**Figure S9.** $T_{FH}$ cells in SLE patients. Frequency of $T_{FH}$ (**A**) and $T_{PH}$ (**B**) cells in peripheral $CD4^+$ T cells. N = 14 per group compared with *t* tests or Mann-Whitney tests. Mean + SEM, *: P < 0.05. (**D** – **E**) Metabolomic analysis showed a mitochondrial dysfunction in the $T_{FH}$ cells from SLE patients. (**D**) Pathway analysis of untargeted metabolomics of SLE and HC $T_{FH}$ cells (N = 3 each). (**E**) Heatmap of differentially enriched metabolites. Metabolite identification was performed through reference to metabolite standards. Additional metabolites were identified by their enrichment in significant pathways and *m*/*z* match. These putatively annotated metabolites are labeled with asterisks (*) in the heatmap.

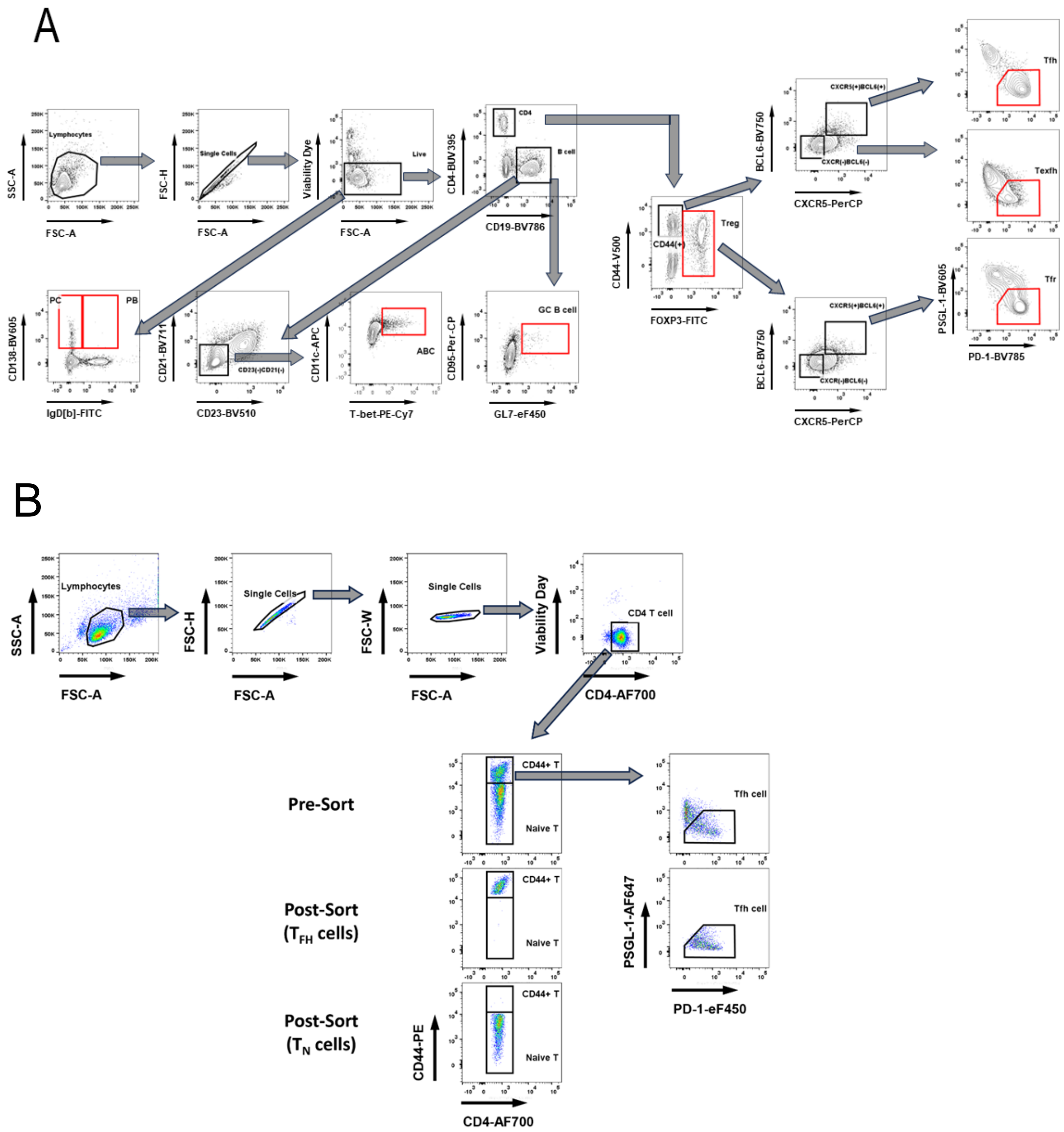


**Figure S10**. (**A**) Gating strategies for mouse CD4$^{+}$ T cells and B cells. (**B**) Representative FACS plots showing sorting strategy and efficiency of mouse $T_N$ and $T_{FH}$ cells. Purified splenic CD4$^{+}$ T cells were sorted into Tn cells as CD44-negative, and $T_{FH}$ cells as CD44$^{+}$ PSGL-1$^{lo}$ PD-1$^{+}$. The pre- and post-sort gates are shown for both cell types.

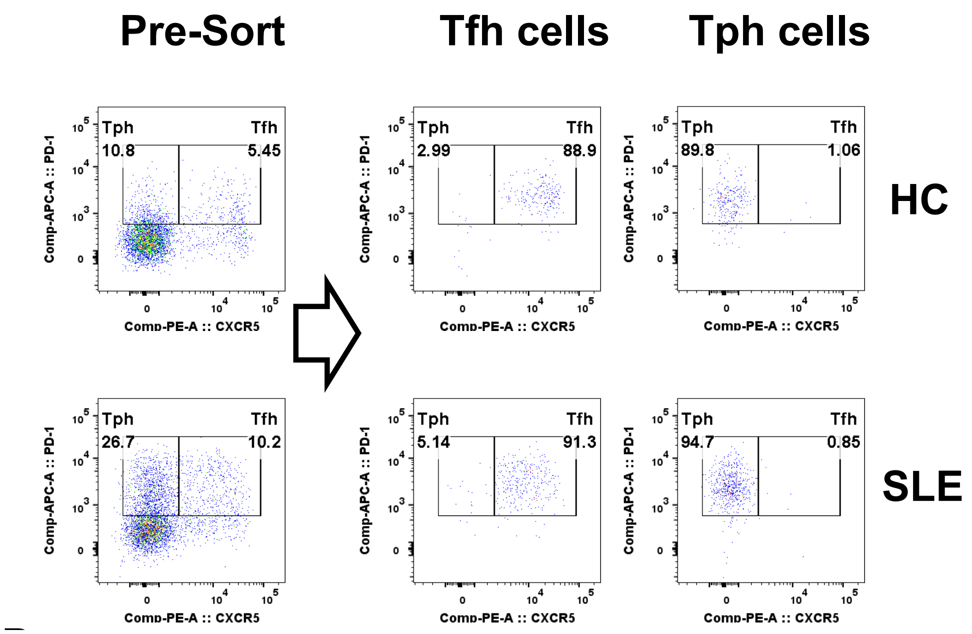


**Figure S11**. Gating and sorting strategy for human $T_{FH}$ and $T_{PH}$ cells from $CD4^+$ T cells
